# Charge-screening dominates signal transduction in electrochemical nucleic acid sensors for aminoglycosides

**DOI:** 10.64898/2026.09.23.753905

**Authors:** Steven Yee, Grace M. Maddocks, H. Tom Soh

## Abstract

Electrochemical aptamer-based (EAB) sensors have emerged as a promising framework for reagentless, real-time molecular monitoring in complex biological matrices. Signal transduction in a prominent benchmark system—targeting aminoglycoside antibiotics—has been nearly universally attributed to target-specific, binding-induced conformational changes that modulate the distance between a redox reporter and the electrode surface. Here, using a suite of interfacial electrochemical and optical characterization methods, we systematically show that charge-screening, not structure switching, dominates the signaling mechanism in the case of the aminoglycoside EAB sensor. First, we demonstrate that the legacy aminoglycoside sensor sequence exhibits apparent affinities and relative electron kinetic modulations similar to unrelated control sequences, including generic stem-loop hairpins, unstructured poly-T sequences, and short duplexes. Multi-modal validation via bio-layer interferometry (BLI), circular dichroism (CD), and fluorescence spectroscopy confirms that this promiscuous target association occurs across disparate DNA sequences and without detectable conformational reconfiguration. Further investigation via charge-scaling polyamine assays and on-electrode fluorescence imaging demonstrate that the dominant signal transduction mechanism in these sensors is instead driven by macro-scale electrostatic backbone charge-screening.

## Introduction

Since their inception more than twenty years ago, electrochemical aptamer-based (EAB) sensors have been used for reagentless, real-time monitoring of diverse molecular targets, even in undiluted biological matrices [1–8]. EAB sensors employ electrode-tethered aptamer affinity reagents modified with a redox reporter that engages in electron transfer with the electrode under electrochemical interrogation. In principle, ligand binding shifts the equilibrium of surface-bound aptamers to alter the ensemble-averaged electron transfer rate—a mechanism conventionally attributed to target-induced changes in aptamer conformation [2, 5, 7, 9]. EABs combine the generalizability of affinity-based approaches with the convenience of reagentless detection and electronic integration. The continued maturation of this technology holds promise for deepening our understanding of pharmacokinetics via real-time sensors and for the realization of personalized medicine through continuous monitoring of diverse biomarkers using point-of-care, wearable, or implantable systems.

One particular EAB reported to specifically bind aminoglycoside antibiotics has featured prominently in the scientific literature since its initial demonstration in 2010 [10]. Since then, this sequence continues to be widely utilized to showcase differing electrochemical interrogation techniques [11, 12] or *in vivo* deployment strategies aimed at improving sensor convenience, sensitivity, or longevity [2, 13–22]. These reports have typically attributed changes in electron transfer rate to specific binding-induced or - stabilized changes in the aptamer’s fold or conformation [2, 10–13, 21]. Other distinct mechanisms have been posited to underlie electrochemical transduction in particular EAB sensors. For example, the signaling mechanism of the three-way junction cocaine/quinine-sensing aptamer was demonstrated to rely on an aptamer-methylene blue interaction that is competitively disrupted by target binding [23]. DNA-mediated charge transport has also been heavily debated as a transduction mechanism for double-stranded DNA constructs either conjugated to or intercalated by redox active molecules [24–27]. However, the prominence of this mechanism in the EAB context is likely limited by the longer alkane linkers typically used in EAB reporter attachment, creating conditions that probably favor the collision-based mechanism for most EABs. Recent literature has reported confounding effects during aminoglycoside detection with EAB sensor controls [28], suggesting that DNA might undergo non-specific structural changes upon aminoglycoside binding or that aminoglycoside-induced changes in ionic strength drive sensor promiscuity. These confounding effects are further informed by prior studies on polycation-induced DNA condensation, where backbone charge neutralization plays a major role in reducing DNA solubility [29, 30]. These reports prompted us to investigate the degree to which the conventional structure-switching aptamer model explains signaling in the aminoglycoside EAB.

Herein, we employ a suite of biophysical techniques—including bio-layer interferometry (BLI), fluorescence spectroscopy, circular dichroism (CD) spectroscopy, and electrochemistry—to demonstrate that the benchmark aminoglycoside DNA sensor does not operate via target-specific, conformationally-driven transduction. Instead, signal change arises from non-specific, promiscuous interactions between aminoglycosides and nucleic acid structures. Indeed, we demonstrate that a wide range of methylene blue (MB)-tagged DNA sequences exhibit sensitivities to aminoglycoside targets that are comparable to that of the reported DNA aminoglycoside aptamer. We hypothesize that signal transduction in these systems results primarily from a charge-screening effect, in which the promiscuous and sequence-independent binding of cationic aminoglycosides to negatively-charged DNA reduces repulsion from the electrode surface, thereby increasing collision efficiency. This effect is exacerbated under the negative potentials necessary for interrogation of methylene blue. We validate this hypothesis through various assays that confirm sequence-agnostic binding, a lack of structural change in electrochemically-signaling constructs, a similar electrochemical response to other polycationic molecules that are known to interact promiscuously with the DNA backbone, and target-induced DNA approach to the underlying electrode.

## Results and Discussion

### Origins of the aminoglycoside DNA EAB

The original aminoglycoside aptamer, first reported in 1995 by Wang et al., is an RNA sequence that reportedly binds tobramycin with a sub-nanomolar dissociation constant (*K*_*D*_) as determined by a fluorescence competition assay [31, 32]. This sequence has a simple predicted hairpin structure in the absence of target, and NMR analysis indicates that the stem of this structure exhibits minimal disruption when bound to tobramycin [33]. The DNA homolog of this aptamer was introduced by Rowe *et al*. in 2010 as an EAB, with terminal thiol and MB modifications for electrochemical interrogation [10]. The DNA homolog EAB was intended to extend sensor longevity in biological matrices, but exhibited greatly reduced apparent affinity (EC_50_ = 1.4 mM) relative to the corresponding RNA EAB (EC_50_ = 350 μM) [10]. Sensitivities of both EABs were several orders of magnitude worse than the 0.77 nM *K*_*D*_ originally reported for the RNA aptamer. Notably, the original report by Rowe *et al*. indicated that the DNA homolog exhibited a negligible CD shift upon tobramycin association, suggesting no meaningful structural change in response to target binding. This is in contrast with the original RNA aptamer, for which a CD shift is attributed to changes in the aptamer loop upon binding.

The authors assessed the specificity of the DNA EAB relative to a control sequence based on an IgGE-binding DNA aptamer using square-wave voltammetry (SWV). They reported a decrease in peak current in the aminoglycoside EAB but not in the control, concluding that target binding specifically triggers a decrease in effective electron transfer rate in the aminoglycoside EAB. However, the authors only assessed SWV response at a single frequency, masking inadequacies in the negative control experiment. An EAB’s SWV titration response at a single frequency is insufficient to determine whether the sensor’s average electron transfer rate is accelerating or decelerating, because a single frequency response reflects only relative changes in the number of electron transfer events at the corresponding time scale. By contrast, comparing multiple SWV frequencies across a target titration indicates the direction of average electron transfer rate change. Indeed, the authors’ conclusions were subsequently contradicted by multiple reports using multi-frequency SWV analysis or chronoamperometry that clearly demonstrated an *increase—*rather than a decrease—in electron transfer rate for this EAB with rising aminoglycoside concentrations [11, 12]. Nevertheless, the EAB research community has continued to employ this DNA sequence for aminoglycoside sensing and attribute its signaling mechanism to aptamer conformation despite the lack of structural evidence for binding-induced hairpin disruption or stabilization and the poor sensitivity of the DNA EAB [11–18, 21, 22]. This disconnect prompted us to reevaluate this widely-used sensor in an effort to better understand the signaling mechanism at play.

### Sequence-agnostic electrochemical response to aminoglycosides across diverse DNA-MB controls

We first sought to resolve the conflicting reports pertaining to aptamer specificity and target promiscuity. To do this, we compared the electrochemical response of the canonical aminoglycoside aptamer to various unrelated control sequences including the original IgGE-binding control sequence (a one-base offset hairpin) used by Rowe *et al*., a random stem-loop hairpin, and a completely unstructured poly-T_26_ sequence (**Fig. 1**). Each construct was modified with a terminal thiol tether and a distal MB redox reporter. If signal transduction is driven by a target-specific conformational transition unique to the aminoglycoside aptamer, the control sequences should exhibit a minimal electrochemical response to tobramycin.

**Figure 1.**
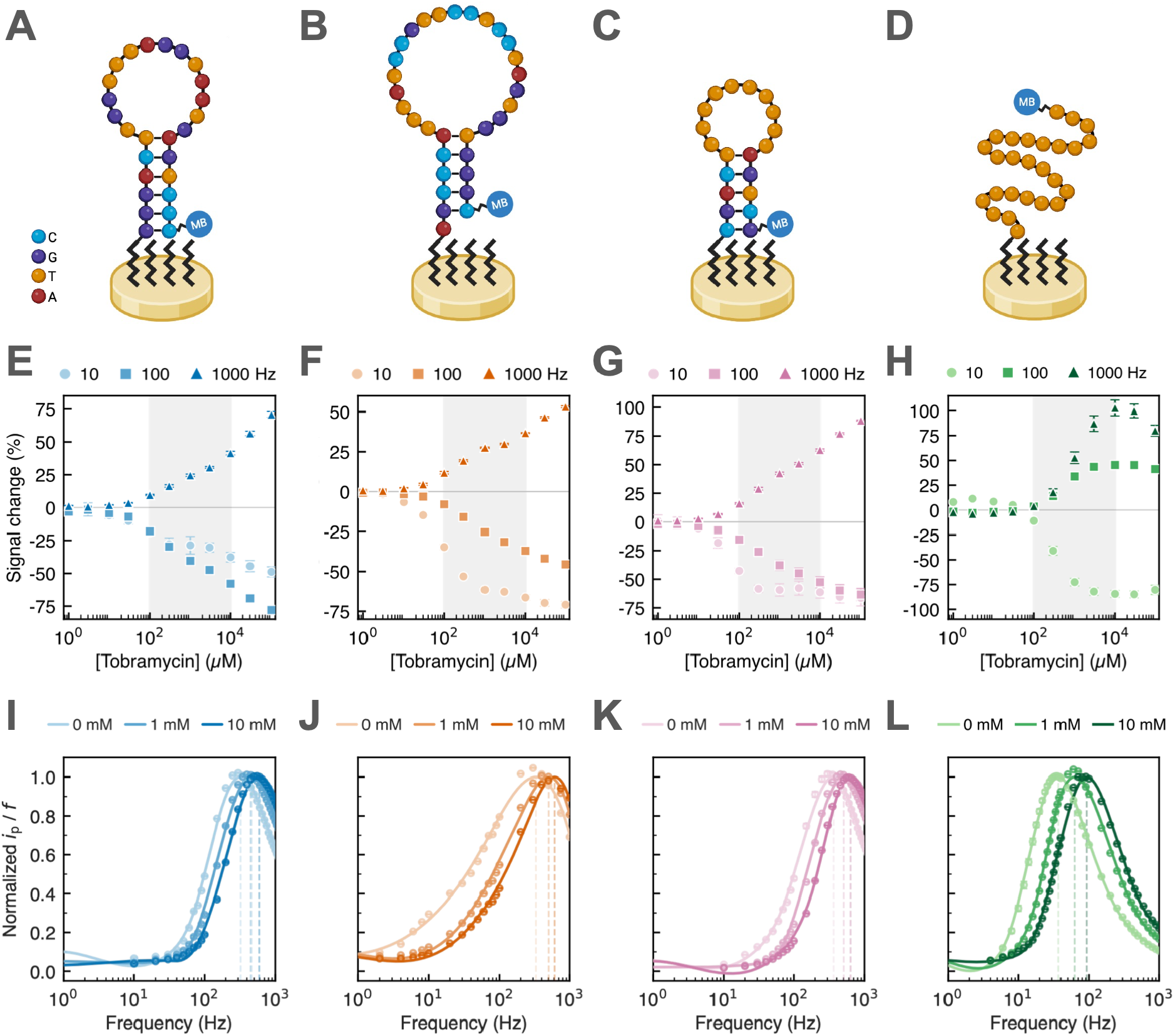
The electrochemical response of DNA-MB sensors to aminoglycosides is not sequence specific. (**A–D**) Illustrative representations of predicted unbound secondary structures for (**A**) the canonical aminoglycoside aptamer, (**B**) the IgGE DNA aptamer utilized as a negative control in [10], (**C**) a random hairpin sequence, and (**D**) a poly-T_26_ control sequence. (**E–H**) Square-wave voltammetry (SWV) signal response curves and (**I–L**) plots of frequency-normalized peak current across frequency for (**E, I**) the canonical aminoglycoside aptamer, (**F, J**) the IgGE aptamer, (**G, K**) the random hairpin control, and (**H, L**) the poly-T_26_ control. All sequences were modified distally with methylene blue (MB). Solid lines in **I–L** indicate spline fits to frequency-normalized current peaks. Dashed lines show the maxima of the frequency-normalized current peaks, which are used as proxies for ensemble-averaged electron transfer rates [34–36]. Error bars in **E–H** represent one standard deviation of three replicates. Error bars in **I–L** represent one standard error on the mean of three replicates.

Strikingly, multi-frequency SWV interrogation across tobramycin concentrations revealed nearly identical apparent affinities across all tested constructs, regardless of sequence (**Fig. 1E–H**). When evaluated using the Mirčeski method (normalizing the peak currents by the applied SWV frequency across a frequency spectrum) to estimate the ensemble-averaged electron transfer rates [34–36], the qualitative behavior of the control sequences closely mirrored that of the canonical aminoglycoside aptamer. The relative changes in observed electron transfer rates were highly similar between the IgGE negative control and the canonical aminoglycoside sensor (**Fig. 1I, J**), with frequency maxima that were shifted only slightly lower for the IgGE control. This is likely due to a minor variation in the baseline electron transfer rate, consistent with the known one-base offset in the hairpin stem of the IgGE control sequence. For all sequences, tobramycin titrations yielded apparent binding curves with similar profiles, demonstrating that the observed electrochemical signaling is largely independent of the nucleic acid sequence employed. We also tested these sequences against kanamycin, a closely-related aminoglycoside for which the original aptamer was reported to exhibit cross-reactivity. We observed similar sequence-agnostic responses, although the magnitude of the poly-T_26_ control response to kanamycin was reduced compared to tobramycin and to the responses of the other sequences to either target (**Fig. S1**).

To ensure that these target-induced equilibrium shifts were not driven by global changes in our testing buffer, we performed a series of controls against pH and ionic strength effects that could affect our reporter or generically impact DNA behavior. Because the electron transfer rate of methylene blue is known to change with pH [37–40], we measured the pH of our Tris-based testing buffer across a range of tobramycin sulfate concentrations from 0 to 100 mM and observed a slight downward shift from pH 7.6 to 7.48 between 100 μM and 10 mM, returning to pH 7.63 at 100 mM. In contrast, increasing concentrations of kanamycin monotonically modulated our buffer pH from 7.6 up to 8.15 (**Fig. S2**). When we fixed the solution pH at 8.15 across target concentrations, we observed a virtually identical kanamycin response to that obtained in the original buffer (**Fig. S3**). For both analytes, the apparent electron transfer rates increased for all sequences with increasing target concentration despite opposing trends in target-induced pH change. Because DNA behavior is dependent on ionic strength [9], we additionally controlled for the concomitant increase in sulfate that accompanies tobramycin and kanamycin when utilized in their sulfate salt form in these experiments. When we titrated sodium sulfate against MB-DNA immobilized electrodes, isolating the impact of bulk ionic strength and salt addition, we observed no significant signal changes (**Fig. S4**). We also assessed the potential role of redox reporter-ligand competition as a potential signaling mechanism by observing MB-DNA fluorescence in solution across a kanamycin titration and did not find strong fluorescence recovery to support this hypothesis (**Fig. S5)**.

The fact that we consistently observe a robust aminoglycoside response across many disparate sequences strongly indicates that this signaling is not sequence- or structure-specific, but is instead the product of a macroscopic interaction between the cationic aminoglycoside and the polyanionic DNA backbone. Crucially, this proposed mechanism of signaling would not require a particular aptamer fold or conformation. This model is further supported by the concentration regime required to elicit a sensor response; across all tested sequences, the apparent half-maximal effective concentration (EC_50_) consistently fell within the hundred micromolar to millimolar range (0.1–1 mM). This result is consistent with previous reports using the aminoglycoside EAB [11, 12, 20, 21], and stands in contrast to the micromolar sensitivities typical of EABs [41–43].

### Label location-agnostic DNA-MB response to aminoglycosides

These findings raised the question of whether transduction involves any conformational reconfiguration at all. To answer this, we synthesized a variant of the canonical aminoglycoside sensor in which we replaced the standard distal 3′ MB label on the DNA sequence with an internal MB modification positioned immediately adjacent to the 5′ thiol tether (**Fig. 2A**). If the observed signal results from a binding-induced structural transition in the aptamer fold, this response should be eliminated by trapping the reporter at a fixed, proximal position next to the gold surface.

**Figure 2.**
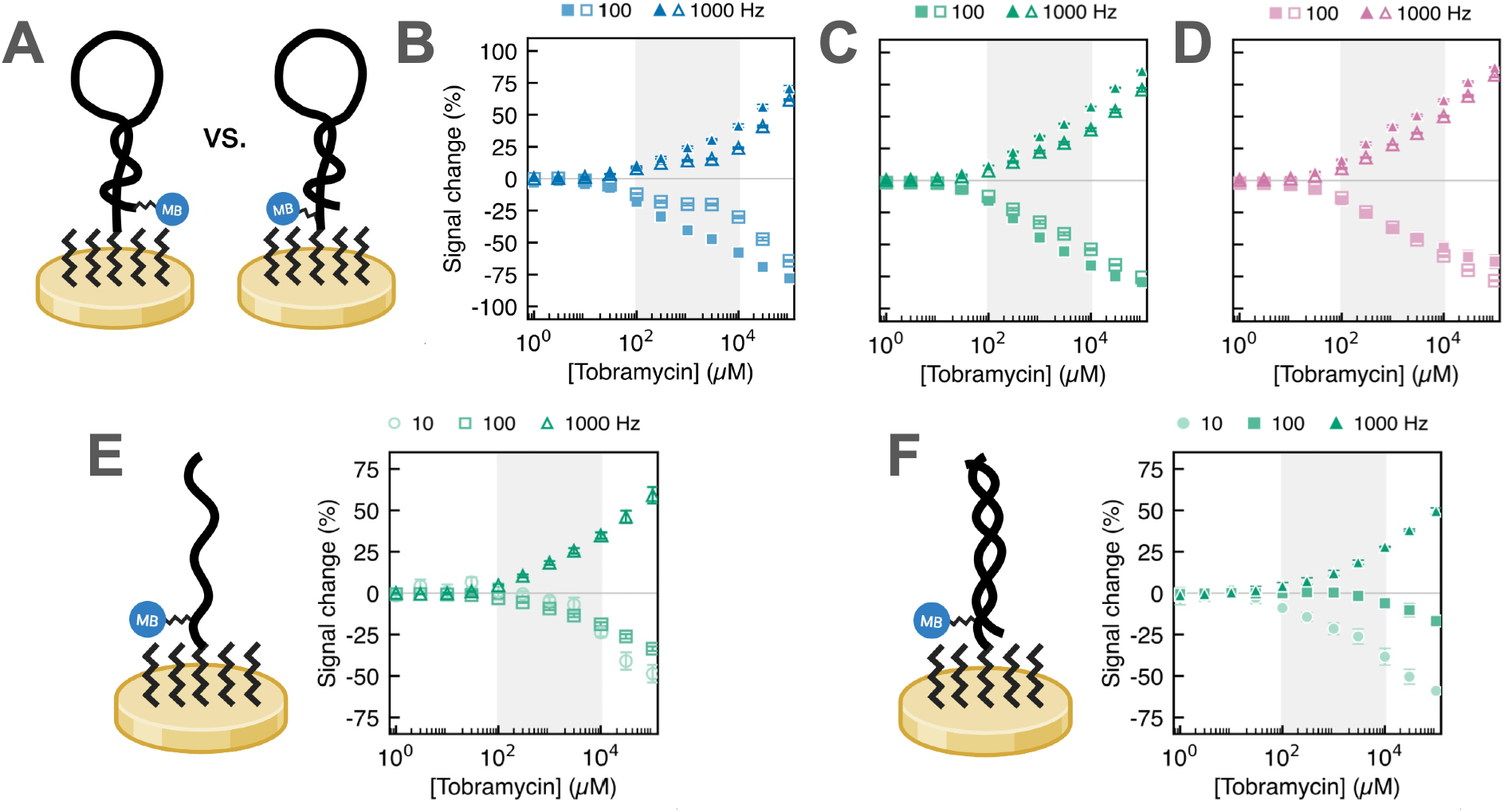
The electrochemical response of DNA-MB sensors to aminoglycosides does not require conformational rearrangement. (**A**) Comparing a conventional, distally-labeled MB (left) to an alternative sensor design (right) in which the MB is labeled immediately proximal to the thiol anchor, restricting its movement. (**B–D**) Electrochemical response of distally-labeled (solid markers) and proximally-labeled (hollow markers) sensors as determined by SWV for (**B**) the canonical aminoglycoside sequence and (**C, D**) two random hairpin-forming sequences. (**E, F**) Evaluating the response of single-stranded DNA with a proximal MB label to tobramycin (**E**) unhybridized or (**F**) hybridized with its full complement, eliminating the possibility of target-induced structure switching as a signaling mechanism. We show here a subset of the interrogation frequencies: 10 Hz (circles), 100 Hz (squares), 1 kHz (triangles). Error bars represent one standard deviation of three replicates.

Instead, we observed that the relative changes in apparent electron transfer rate and signal gain persist regardless of reporter position. The proximally-labeled aminoglycoside sensor and its distally-labeled counterpart exhibited similar tobramycin-induced electrochemical responses (**Fig. 2B**). We also compared proximally- and distally-labeled versions of two stem-loop hairpin control sequences, which also exhibited negligible differences between labeling schemes (**Fig. 2C, D**). Each hairpin featured a 10-nucleotide (nt) poly-T loop with differing base identities in a 5-base-pair (bp) stem region. These results strongly indicate that signal transduction is independent of aptamer conformation-derived changes in relative reporter-to-electrode proximity.

To ensure that lingering conformational dynamics from our stem-loop-structured DNA constructs were not influencing signal transduction, we evaluated a random 12-mer DNA sequence in both its flexible single-stranded (ss) DNA form and when fully hybridized to a complementary strand to form rigid double-stranded (ds) DNA. For these constructs, the internal MB label was positioned on the surface-tethered strand immediately adjacent to the anchor, while the complementary strand consisted of unmodified, native DNA. A concentration-dependent tobramycin response by the dsDNA construct, which lacks a loop or other flexible domains, would rule out binding-induced conformational change as a source of sensor signaling. Strikingly, both the dsDNA duplex and the unhybridized ssDNA exhibited apparent affinities for tobramycin that fell within the same regime as the original aminoglycoside sensor and the random hairpins (**Fig. 2F, G**). When tested against kanamycin, responses by these ssDNA and dsDNA constructs were also similar to those of the canonical sensor and controls (**Fig. S6**). We do observe differences in sensitivity across frequency for dsDNA and ssDNA, reflecting differences in baseline electron transfer rates (**Fig. S7**). These findings led us to hypothesize that aminoglycosides promiscuously bind nucleic acid structures and transduce electrochemical signal not by the target-specific aptamer folding described in the conventional EAB sensor paradigm (**Fig. 3A**), but rather by neutralizing electrostatic repulsion of negatively charged DNA-MB constructs from the electrode surface (**Fig. 3B**). Our model explains why conserved hairpin structures and proximally-labeled DNA-MB sequences experience accelerated electron transfer rates with increasing aminoglycoside concentration even though these sequences already are dominated by folded conformations that situate the redox reporter near the electrode in the target-free state. Our proposed mechanism does not necessitate an aptamer conformational shift; it requires only that the binding of one or more polycationic molecules attenuates electrostatic repulsion from the working electrode surface.

**Figure 3.**
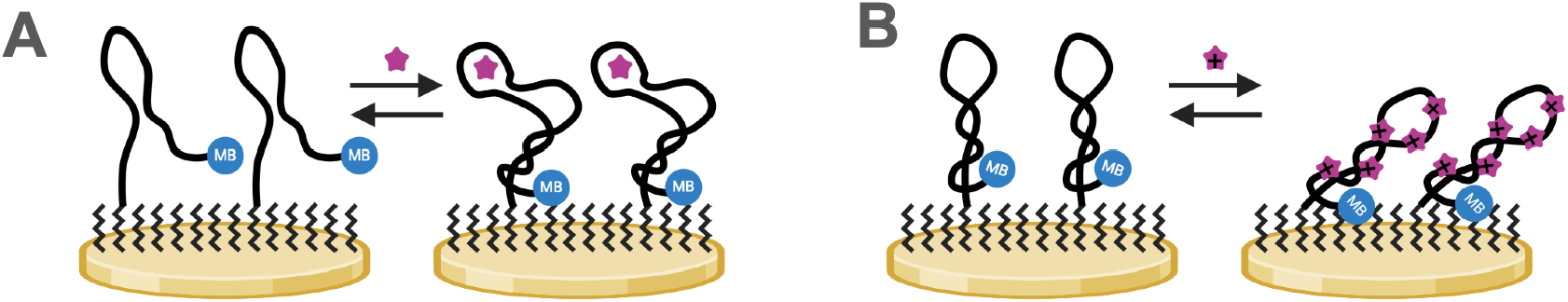
An alternative hypothesis for aminoglycoside EAB signaling. (**A**) The conventional EAB paradigm, in which changes in electron transfer are attributed to changes in the conformation of the DNA. (**B**) Our proposed charge-screening model, in which changes in electron transfer are instead induced by electrostatic screening of the negatively charged DNA backbone by positively-charged targets, reducing the repulsion of the DNA from the surface and increasing the collision rate of the reporter with the surface.

### Non-specific aminoglycoside-DNA association

We first set out to confirm the extent of nonspecific DNA-aminoglycoside interactions by performing bio-layer interferometry (BLI). BLI measures binding interactions based on changes in the refractive index that occur as a result of target association with receptor molecules immobilized on the sensor tip. This optical technique offers an orthogonal approach to our EAB experiments and is not susceptible to electrochemical artifacts.

The BLI results confirm that aminoglycosides interact indiscriminately with nucleic acids. We immobilized onto streptavidin-coated probes a variety of biotinylated DNA constructs and assessed their binding to tobramycin. Our panel included the aminoglycoside aptamer, a scrambled version of the aminoglycoside aptamer, the IgGE aptamer, a random hairpin, and poly-T_26_ sequences. For every tested construct, our reference-subtracted sensorgrams demonstrated a concentration-correlated shift in refractive index indicative of tobramycin association (**Fig. 4** and **Fig. S8**). At each concentration, we subtracted a reference measurement obtained from probes lacking any DNA construct to ensure that each response represents true aminoglycoside-DNA interactions as opposed to non-specific probe interactions. Notably, both association and dissociation evolutions were characterized by an apparently instantaneous response for all sequences, with a slower kinetic tail observed to varying degrees across sequences. The canonical aptamer exhibited a larger contribution from these slower kinetic components compared to the other sequences, but its response was ultimately still dominated by the nearly instantaneous component shared by all sequences. Given this strongly multi-phasic binding behavior, we did not attempt to extract kinetic rate constants from these data. Similar sensorgrams were observed in response to kanamycin (**Fig. S9**) and for scrambled aminoglycoside aptamer or alternative hairpin sequences (**Fig. S8**). The rapidity of the aminoglycoside-DNA association implied by BLI was further corroborated by SWV across sequences and targets (**Fig. S10**).

**Figure 4.**
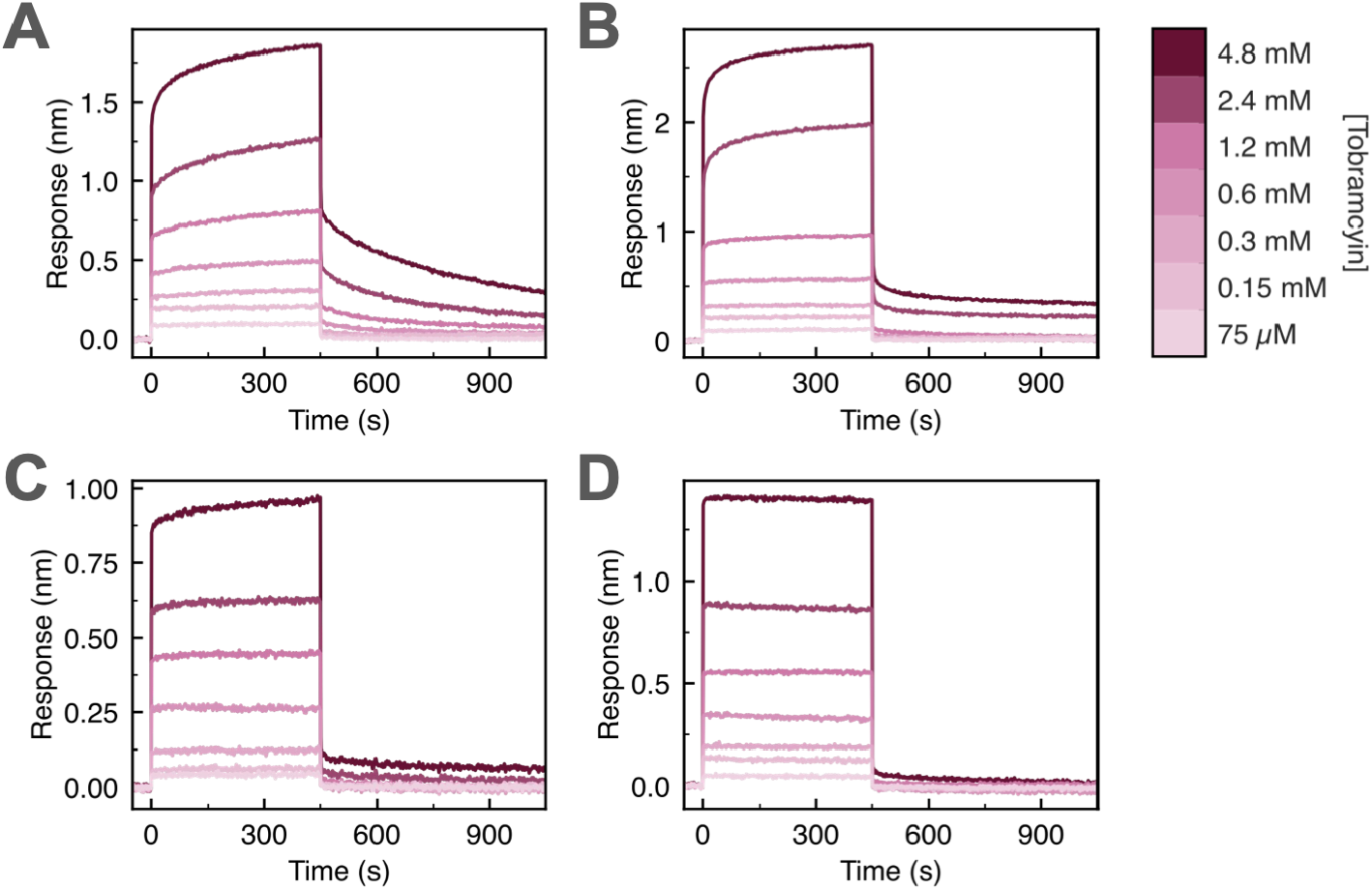
Bio-layer interferometry (BLI) demonstrates tobramycin-DNA association is not sequence specific. (**A–D**) Reference-subtracted BLI sensorgrams for tobramycin against biotinylated variants of (**A**) the aminoglycoside DNA aptamer, (**B**) the IgGE aptamer, (**C**) a stem-loop hairpin, and (**D**) a poly-T_26_ control. Association occurs from *t* = 0 to *t* = 450 s. Dissociation occurs for *t* > 450 s.

### Conservation of DNA structure when bound to aminoglycosides

We next sought to orthogonally verify that aminoglycoside binding does not induce macromolecular conformational changes in various DNA sequences by performing CD spectroscopy. Any meaningful target-stabilized conformational change associated with aminoglycoside exposure should give rise to a distinct shift or redistribution in the characteristic DNA helicity profiles. We tested the canonical aminoglycoside sensor, a random stem-loop hairpin, and a poly-T_26_ sequence and consistently observed a negligible CD spectrum shift upon introduction of tobramycin or kanamycin, even at target concentrations well above the sensor’s electrochemical response threshold (**Fig. 5A–F**). This result is consistent with the original EAB report by Rowe *et al*. [10] and confirms that aminoglycoside-DNA binding elicits no detectable alteration to the underlying nucleic acid structures. For comparison, we also evaluated known structure-switching aptamers reported in the literature for glucose and vancomycin [42, 44], and observed clear target-associated shifts in their CD spectra (**Fig. S11**). Numerical comparisons of relative structural shifts based on these CD spectra are shown in **Table S1**.

**Figure 5.**
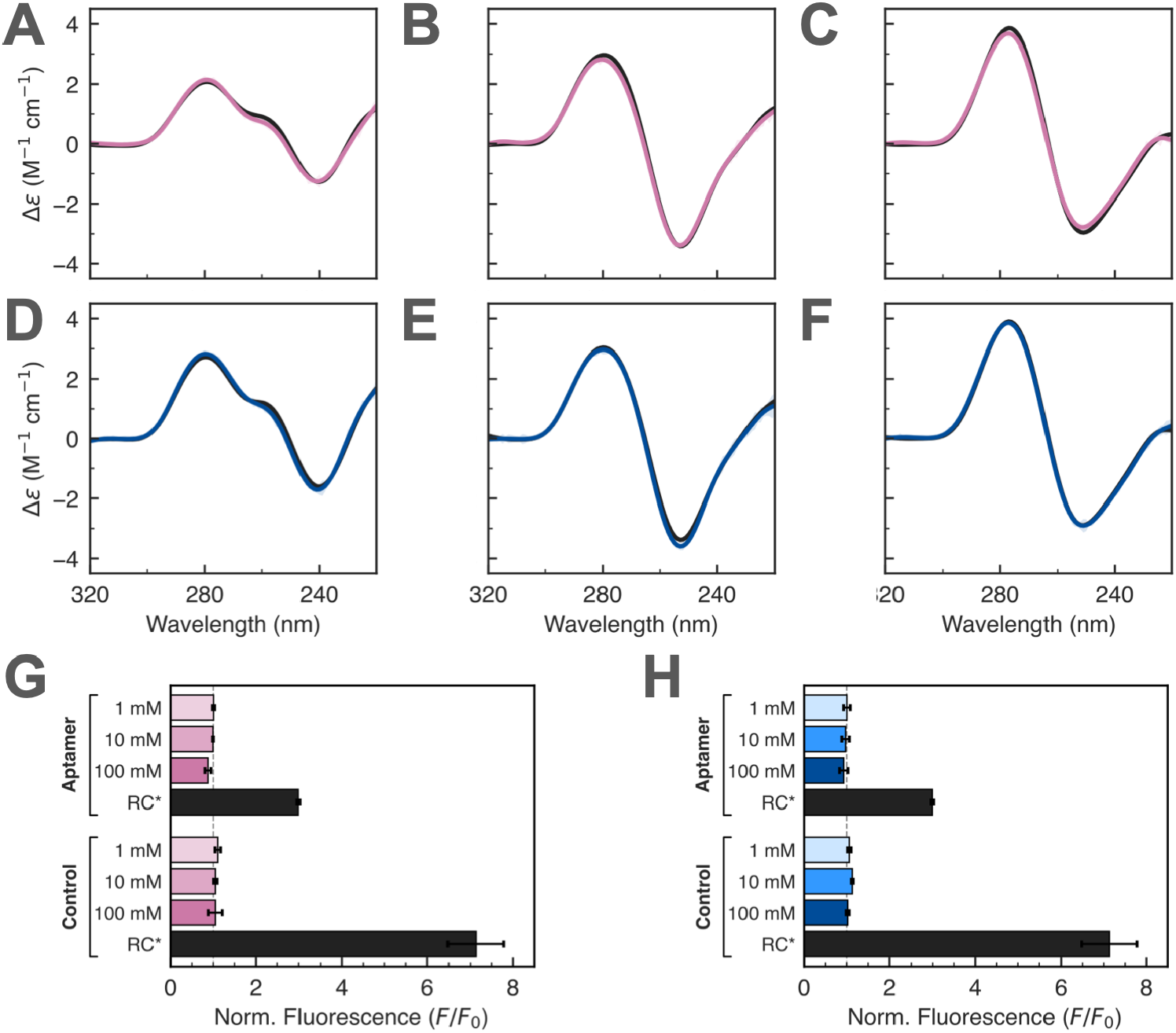
Circular dichroism (CD) and fluorescence indicate that no DNA conformational change occurs in response to aminoglycosides. (**A–F**) CD spectra in the absence (black) or presence (color) of (**A–C**) 100 mM tobramycin or (**D–F**) 100 mM kanamycin for (**A, D**) the canonical aminoglycoside aptamer, (**B, E**) a random hairpin, and (**C, F**) poly-T_26_ sequences. Quantitative relative shifts are available in Table S1. Semi-transparent patches indicate error-propagated single standard deviations. (**G, H**) Fluorescence changes for the canonical aminoglycoside aptamer and random hairpin control variants terminally labeled with fluorophore and quencher pairs in response to tobramycin (**G**) and kanamycin (**H**). For comparison, we also evaluated the response of these sequences to excess quantities of unlabeled reverse complement sequences (RC*) as a positive control. Error bars represent one standard deviation of three replicates.

To further substantiate a lack of structural change, we evaluated versions of the canonical aminoglycoside sensor and the random stem-loop hairpin modified with a dye-quencher pair (Alexa Fluor 594 and Black Hole Quencher-2) at their 3′ and 5′ ends. As both constructs are expected to be dominated by hairpin structures under the ionic conditions tested (**Fig. S12**), we expected to see low fluorescence emission in the target-free state; a sequence undergoing target-induced conformational change would thus be indicated by a target-dependent increase in fluorescence. However, we saw negligible changes in fluorescence for each of these constructs in response to tobramycin or kanamycin titrations (**Fig. 5G, H**). In contrast, we saw multi-fold fluorescence increases when these constructs were incubated with an excess of their respective, unlabeled reverse complements. These increases further confirm the predominance of folded hairpin structures in the target-free condition for each of these structures. These results strongly indicate that the mechanism for electrochemical response of MB-DNA to aminoglycosides does not involve a large-scale conformational rearrangement.

### Electron transfer rate accelerations for MB-DNA constructs across polycationic targets

Having established that signal transduction is not driven by target-specific conformational changes, we hypothesized that this response may instead arise from the promiscuous accumulation of cationic species along polyanionic DNA backbones. To test this, we electrochemically interrogated a panel of DNA constructs—the canonical aminoglycoside aptamer, a stem-loop hairpin, and an unstructured poly-T_26_ sequence—against not only the aminoglycosides tobramycin and kanamycin, but also the polycationic molecules spermine and spermidine. These polyamines are known to neutralize DNA phosphate backbone charge and can even induce DNA precipitation [30]. We reasoned that if signaling is driven by non-specific electrostatic association, we would observe responses to all of these polycationic targets across all constructs.

In line with our hypothesis, we observed accelerated electron transfer for each DNA construct against all tested targets via the Mirčeski method. We assessed electron transfer rate changes by comparing *f*_peak_ across target titrations relative to a zero target baseline *f*_peak,0_. For each construct, we observed that *f*_peak_ begins to accelerate at similar concentrations, supporting a model in which sensor signals previously attributed to specific aminoglycoside-DNA induced conformational rearrangements instead arise from macroscale charge-screening effects (**Fig. 6C–F**). Our observation of promiscuous association across polycationic targets is independently supported by BLI measurements, which verifies that spermine interacts non-specifically with these disparate DNA constructs (**Fig. S13**). Notably, when we compare electrochemical responses within target classes (*i*.*e*., aminoglycosides or polyamines), the more highly charged targets—tobramycin and spermine—exhibited earlier onset of electron transfer rate acceleration compared to their less positively-charged analogues, kanamycin and spermidine. This relative behavior further corroborates our charge-screening model, suggesting that when two targets are structurally similar but differ in net charge, additional positive charge can increase the target’s efficacy in screening the DNA backbone.

**Figure 6.**
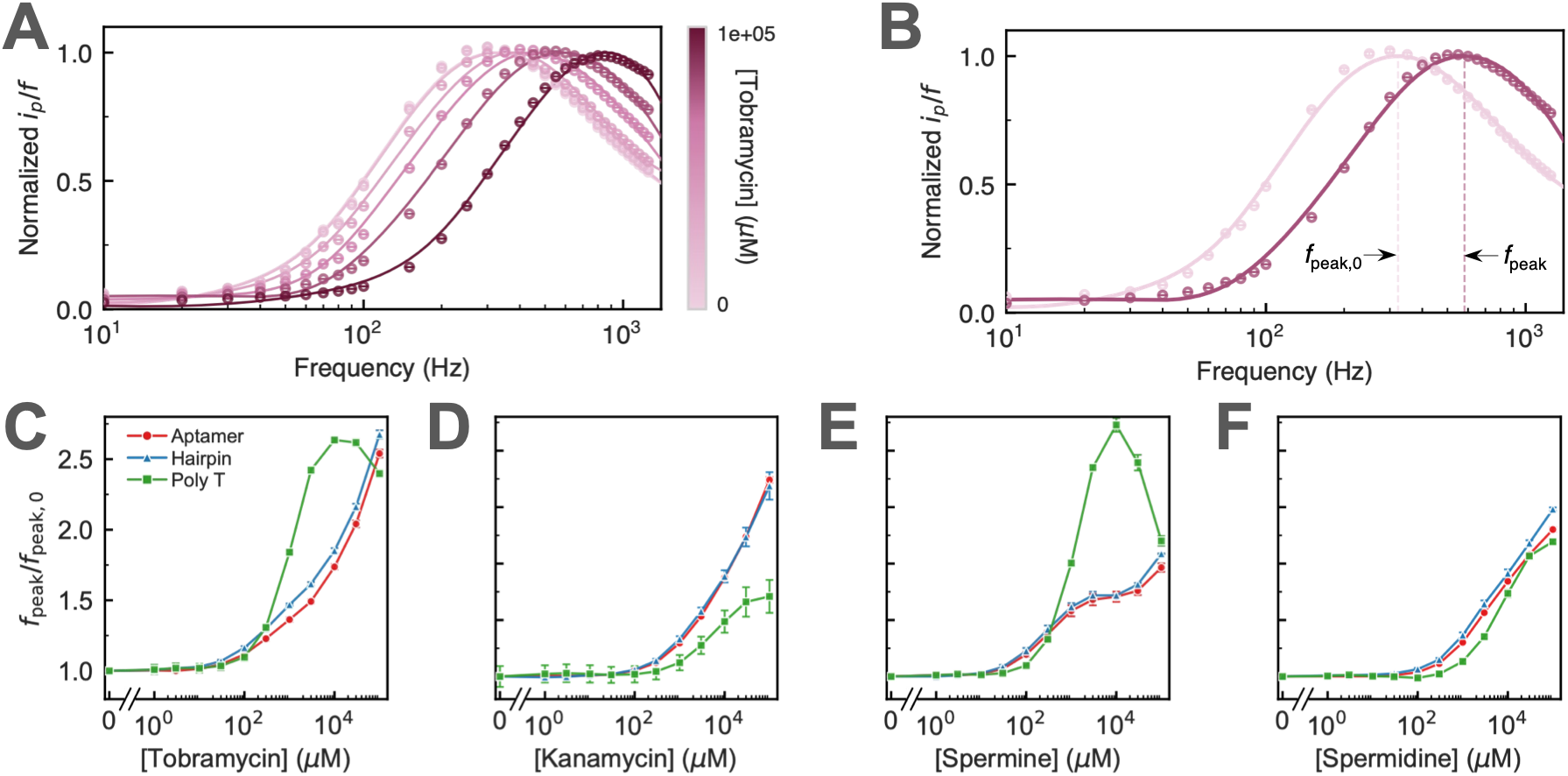
Multi-frequency SWV analysis exhibits similar trends in apparent electron transfer rate across polycationic targets. (**A**) A representative Mirčeski analysis of the canonical aminoglycoside EAB against tobramycin and (**B**) an annotated Mirčeski analysis indicating peak frequency for the baseline and target-bound conditions (*f*_peak,0_ and *f*_peak_, respectively). (**C–F**) Plots of normalized peak frequency (*f*_peak_ / *f*_peak,0_) for the canonical aminoglycoside aptamer, a hairpin control, and a poly-T_26_ control across titrations of (**C**) tobramycin, (**D**) kanamycin, (**E**) spermine, and (**F**) spermidine, respectively. Error bars represent standard error on the mean of three replicates.

Although the general response to polycationic targets was an increase in electron transfer rate, we observe some structure-dependent nuances. The two hairpin forming sequences—the aminoglycoside aptamer and the stem-loop control—behaved nearly identically across target titrations, but the behavior of the unstructured poly-T_26_ diverged in some respects. For instance, we observed for the poly-T_26_ a more rapid acceleration of electron transfer at low concentrations of tobramycin and spermine, followed by a deceleration at concentrations greater than 10 mM (**Fig. 6C, E**). At high concentrations of spermine, some deceleration was observed for all sequences, an effect we attribute to spermine’s aggressive reduction of pH (**Fig. S2**). The poly-T_26_ control also showed a weaker response to kanamycin than the other sequences (**Fig. 6D**), while its acceleration rate for all other targets was consistently greater than or equal to that of the canonical aptamer and hairpin control. These findings indicate that while disparate DNA-MB constructs share broadly similar electrochemical responses when exposed to polycations, structure-dependent nuances remain.

### Target-induced collapse and reduction of electrophoretic modulation of tethered constructs

To directly visualize how this macroscopic charge-screening modulates the physical orientation and mechanical dynamics of the DNA ensemble under an applied voltage, we performed *in situ* fluorescence imaging. For these experiments, we replaced the MB at the 3′ terminus of the canonical aminoglycoside aptamer with a sulfo-Cy3 fluorophore, for which the gold electrode serves as a distance-dependent quencher [45–49]. We expected that the negative potentials used for MB SWV interrogation (−0.45 V to −0.1 V vs. Ag pseudo-reference) would drive the polyanionic DNA away from the surface due to electrostatic repulsion, resulting in an increase in fluorescence during interrogation (**Fig. 7A**). Our model predicts that the association of cationic aminoglycosides will screen the DNA backbone and attenuate this repulsion, leading to a concentration-dependent reduction in fluorescence intensity and relative modulation under voltage.

**Figure 7.**
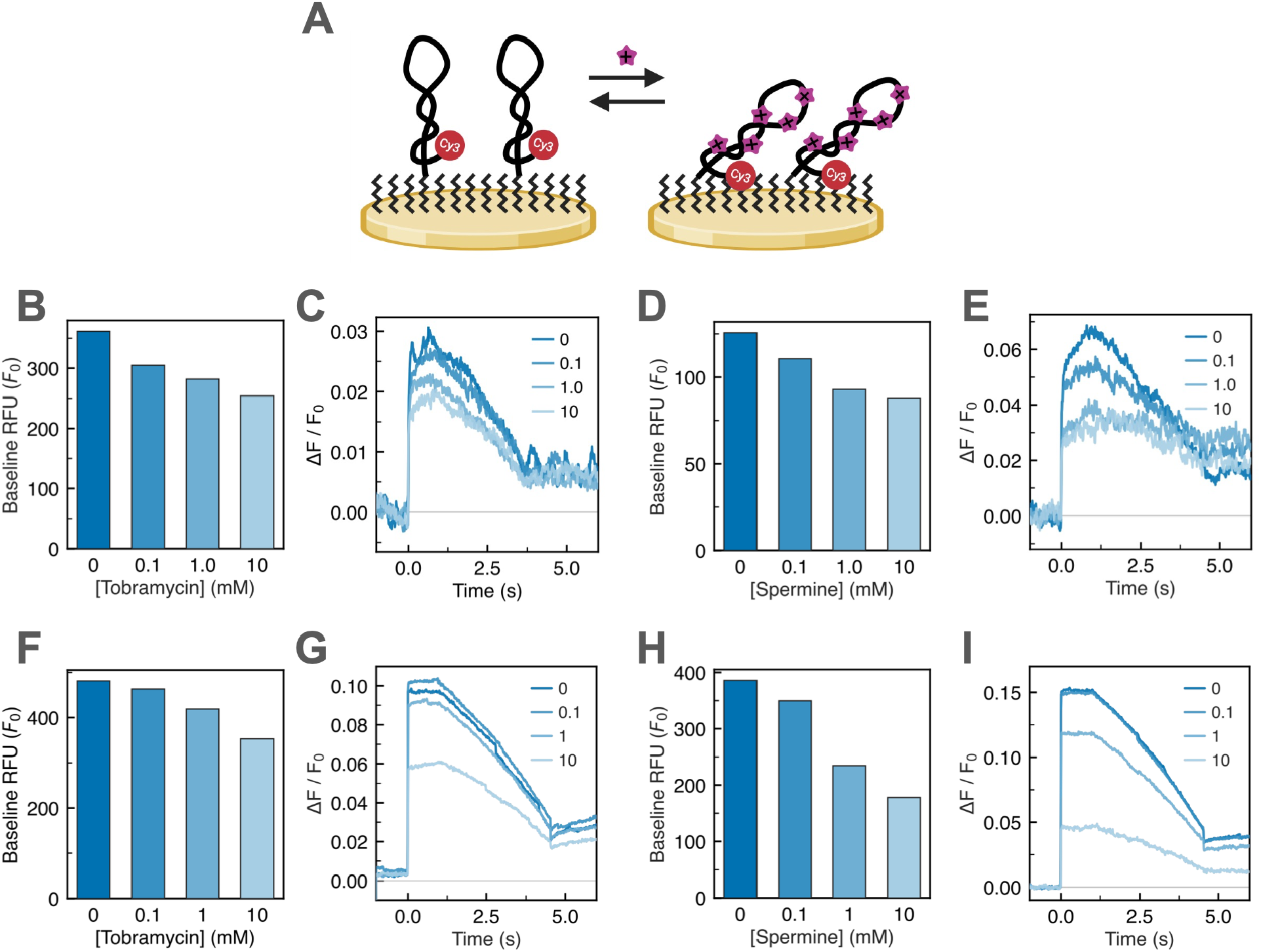
Imaging of fluorophore-modified DNA on an electrode surface supports the charge-screening hypothesis. (**A**) To optically investigate our charge-screening hypothesis, the MB labels were swapped for Cy3 or Cy3B. The average proximity of the label to the electrode surface is transduced as fluorescence, as gold has a distance-dependent quenching effect on these fluorophores. (**B**) Baseline fluorescence intensity of Cy3-labeled aminoglycoside aptamer before SWV voltage interrogation at 0 V vs. Ag pseudo-reference and (**C**) relative fluorescence intensity modulation of the Cy3-labeled aminoglycoside aptamer under an applied SWV waveform at 100 Hz at increasing tobramycin concentrations. (**D**) Baseline fluorescence and (**E**) relative intensity modulation of Cy3-labeled aminoglycoside aptamer at increasing spermine concentrations. (**F**) Baseline fluorescence and (**G**) relative intensity modulation of a poly-T_26_-Cy3B construct at increasing tobramycin concentrations. (**H**) Baseline fluorescence and (**I**) relative intensity modulation of a poly-T_26_-Cy3B construct at increasing spermine concentrations. All concentrations are millimolar.

Our optical measurements confirmed this prediction. Both the baseline fluorescence prior to waveform application (electrode potential at 0 V) and the relative modulation in fluorescence amplitude during waveform application (square-wave ramp from −0.45 V to −0.1 V) decreased monotonically with target concentration. This trend was consistent across the polycation panel—including tobramycin (**Fig. 7B, C**), kanamycin (**Fig. S14**), and spermine (**Fig. 7D, E**). This charge-screening effect also proved reversible for aminoglycosides when we washed the sensor with target-free buffer, although we observed some hysteresis—likely resulting from incomplete washing and photobleaching effects (**Fig. S15A, S15B**). Testing with spermine showed poor reversibility after the highest tested concentration of 10 mM (**Fig. S15C**), a result corroborated by the limited dissociation observed with BLI (**Fig. S13**). Titrations of these targets against the same constructs in solution (**Fig. S16**) do not explain the quenching effects observed in these on-electrode experiments. These data provide direct optical corroboration of our electrostatic model. While the use of a sulfonated dye introduces a negative charge at the reporter site, the global electrostatic responsiveness of the DNA is overwhelmingly dictated by the polyanionic DNA backbone, which features a charge density that is orders of magnitude larger than a single fluorophore modification. To further eliminate the possibility of interference by charged dyes, we also imaged the poly-T_26_ control modified with Cy3B, a fluorophore that is neutrally charged at the pH levels applicable herein. As with the Cy3-labeled aminoglycoside aptamer, we observed that both tobramycin and spermine produced a concentration-dependent reduction in baseline fluorescence and relative voltage-induced fluorescence modulation (**Fig. 7F–I**). Interestingly, kanamycin titration had a weaker effect on the fluorescent signal (**Fig. S17**), consistent with our observation of a reduced electrochemical response magnitude for the poly-T_26_-MB to kanamycin. We were unable to successfully image the aminoglycoside aptamer with a Cy3B modification due to insufficient fluorescence intensity under identical immobilization procedures. We attribute this low fluorescence to the enhanced quenching of Cy3B by the gold electrode relative to Cy3, as well as the hairpin conformation of the aptamer relative to the unstructured poly-T_26_ sequence. However, the fact that we observed the same pattern of fluorescence modulation with the neutral Cy3B on the poly-T_26_ construct indicates that the charge of the dye is not a dominant factor in these experiments. The similar responses of disparate sequences to both aminoglycosides and known backbone-binding polyamines like spermine strongly suggests that these molecules impact DNA EABs in a highly similar fashion. We therefore conclude that the electrochemical response of MB-modified DNA to aminoglycoside antibiotics is in fact driven by a sequence-independent, electrostatic screening effect rather than sequence- or structure-specific changes in DNA folding.

## Conclusion

The results of our investigation collectively indicate that this widely-used aminoglycoside EAB sensor does not operate via the conventionally assumed mechanism of target-specific intramolecular conformational switching. Instead, signal transduction in this legacy system is dominated by non-specific interactions with polycationic aminoglycosides that result in DNA backbone charge-screening. Although the legacy aminoglycoside DNA sensor sequence does not provide additional sensitivity relative to the macroscale charge-screening effect, we do note two recent reports of aptamers for aminoglycoside detection—one based on a new selection toward kanamycin [50], and another that converts the legacy sequence into a fully modified 2′-OMe analogue [19, 20]. Our initial investigations utilizing circular dichroism do suggest that these sequences undergo structural rearrangement upon target binding (**Fig. S18, Table S1**), and preliminary electrochemical results suggest that EABs based on these aptamers may be capable of aminoglycoside detection below the regime where non-specific charge-screening effects dominate (**Fig. S19**).

To protect future sensor development from being confounded by these non-specific interactions, we recommend that the characterization of new electrochemical nucleic acid sensors routinely incorporate a minimal matrix of sequence and spatial controls. Testing a variety of control sequences of equal length to the aptamer of interest—including completely scrambled or generic poly-T sequences—is an essential step to quickly rule out nonspecific interactions and macro-scale electrostatic charge screening. This test is particularly important when targeting highly-charged analytes at concentrations in the high micromolar to millimolar range. Furthermore, relocating the redox reporter from its traditional distal position to an internal site proximal to the surface anchor offers a simple test to verify whether signal transduction is truly dependent on conformational change, alongside more traditional—but still essential—techniques like CD spectroscopy. Adopting these straightforward verification steps alongside orthogonal biophysical methods will provide a rigorous framework for confirming that newly-designed sensors exhibit true sequence- and target-specific binding and structure-switching capabilities, thereby accelerating the realization of high-fidelity, validated platforms for personalized medicine and *in vivo* molecular monitoring.

## Acknowledgements

The authors acknowledge Michael Eisenstein for editorial feedback and Jingjia Chen and the Huang Lab at Stanford for their assistance with training on the circular dichroism instrument. S.Y. acknowledges funding from the Stanford Graduate Fellowship. G.M. acknowledges funding from the National Science Foundation Graduate Research Fellowship Program (NSF GRFP). This work was supported by the National Institute of Biomedical Imaging and Bioengineering of the National Institutes of Health (R01EB038309), the Leona M. and Harry B. Helmsley Charitable Trust, and the Stanford Center for Digital Health.

## Supporting Information

The Supporting Information is available free of charge.

Materials and methods. Supplemental data for electrochemical, pH, fluorescence, bio-layer interferometry, circular dichroism spectroscopy measurements. Predicted unbound DNA secondary structures. DNA sequences. (PDF)

