## Supporting Information for "Charge-screening dominates signal transduction in electrochemical nucleic acid sensors for aminoglycosides"

for

This file includes:

Materials and Methods

Figures S1–S19

Tables S1 and S2

### Materials and Methods

#### *Materials*

Ultrapure water (10977), 1 M Tris-HCl pH 7.5 (15567), 1× Phosphate Buffer Saline (10010), 10× Phosphate Buffer Saline (BP399), Sodium Bicarbonate (S233), 1 M Magnesium Chloride (AM9530G), Spermidine Trihydrochloride (215100010), Alexa Fluor 594 NHS Ester (A20004), 1 N NaOH (SS266), and HPLC-grade Acetonitrile (A998) were sourced from Thermo Fisher. 1 N HCl (H9892), tris(2-carboxyethyl)phosphine (TCEP) (C4706) and D-(+)-Glucose (G8270) were sourced from Sigma Life Science. 1 M HEPES Buffer (1688449) was sourced from MP Biomedicals (Solon, OH). 5 M Sodium Chloride (BM-244) was sourced from Boston BioProducts, Inc. (Milford, MA). Atto MB2-NHS Ester was sourced from Leica Microsystems (Product # AD-MB2-31; South San Francisco, CA). Cy3-NHS Ester was sourced from Apex Bio (Catalog # A8107; Houston, TX). Cy3B-NHS Ester was sourced from Vector Labs (SKU # FP-1321-1; Newark, CA). Tobramycin Sulfate was sourced from G-Biosciences (Catalog # RC-1122; St. Louis, MO). Kanamycin Sulfate (0408-EU-25G) along with 0.2-μm modified nylon spin filters (82031-356) were sourced from VWR (Radnor, PA). Spermine tetrahydrochloride was sourced from Ambeed (Catalog # A134234; Buffalo Grove, IL). All reagents used during DNA synthesis, as well as 2 M Triethylamine Acetate were purchased from Glen Research (Sterling, VA). Key modifications include biotin (10-1953), C6-disulfide (10-1936), 5'-DMS(O)MT-Amino-Modifier C6 (10-1907), 5'-BHQ-2 (10-5932), Spacer 18 (10-1918), dabcyI-dT (10-1058), 3'-PT-Amino-Modifier C6 CPG (20-2956), 3'-PT-Amino-Modifier C3 CPG (20-2954), 3'BHQ-2 CPG (20-5932), 2'-OMe-A-CE (10-3100), 2'-OMe-Ac-C-CE (10-3115), 2'-OMe-G-CE (10-3121), 2'-OMe-U-CE (10-3130). 500-mL 0.22-μm filter systems were sourced from Corning (Corning, NY). HPLC vials (5182-0716), screw caps (5190-7024), and inserts (5181-1270) were sourced from Agilent (Santa Clara, CA).

#### *DNA synthesis, labeling and purification*

All DNA oligos were synthesized either in-house or by Integrated DNA Technologies (IDT; Coralville, IA; see **Table S2** for sequences). In-house oligos were synthesized on an Applied Biosystems Expedite 8908 nucleic acid synthesis system using phosphoramidite chemistry. Oligos procured from IDT were ordered with standard desalting.

All dye-conjugated oligos were first modified with amino groups during synthesis and then subsequently modified with methylene blue (MB)-NHS Ester, Cy3-NHS Ester, or Cy3B-NHS Ester. NHS-Ester reactions were conducted overnight at 25°C or 4°C in 100 mM sodium bicarbonate with a 15-fold molar excess of dye over DNA. After overnight reaction, samples were desalted with Gel-Pak™ 0.2 Desalting Columns (Glen Research), filtered through a 0.2-μm spin filter (VWR), and purified via HPLC on an Agilent 1260 Infinity II instrument equipped with either an Agilent (Part # PL1512-3301) or Hamilton

(Part # 79425) reversed-phase C18 column. HPLC utilized acetonitrile and 100 mM triethylamine acetate in water as mobile phases. Mobile phases were filtered by a 0.22-μm vacuum filter before use. After HPLC purification, samples were dried down in a Thermo Fisher SRF110 centrifugal vacuum.

##### *Square-wave voltammetry*

Unless otherwise noted, all electrochemical testing was conducted in the buffer from the original report of the aminoglycoside EAB: 20 mM Tris, 100 mM NaCl, 5 mM MgCl<sub>2</sub>, pH 7.5. All testing was conducted on 220BT or C220BT commercial screen-printed electrodes featuring gold-based working and counter electrodes and a silver-based pseudo-reference electrode (Metrohm, Llanera, Asturias, Spain). All electrochemical measurements were performed on a PalmSens 12-channel MultiEmStat4 LR potentiostat (PalmSens BV, Houten, Netherlands). Electrodes were interfaced with the potentiostat using PS-CONN-2MM connectors (PalmSens BV).

SWV measurements were performed in the MultiTrace4 software in “individual mode” from $-0.45$ to $0$ V with a $1$ mV step, $25$ mV amplitude, and a $1$ s equilibration time (voltage held at $-0.45$ V) at the start of the measurement. SWV peak fitting was conducted using the ASWIFT algorithm as described in [51]. Except when otherwise specified, testing was performed on sessile $100$-μL droplet solutions exchanged by wicking with a Kimwipe at the edge of the droplet followed by immediate deposition of a new droplet. Equilibration for greater than $5$ min before measurement was allowed after droplet exchange.

All presented electrochemical response curves represent averages and standard deviations as error bars for three replicates. Mirčeski analysis plots represent normalized averages and standard errors on the mean as error bars for three replicates. Normalization is relative to the apex of a univariate spline fit to the triplicate data for each condition. $f_{\text{peak}}$ is derived from the fitted apex of the spline fit. The univariate spline fit and vertical $f_{\text{peak}}$ dashed indicator are plotted to guide the reader.

##### *Electrode preparation*

All electrodes were cleaned before functionalization using cyclic voltammetry (CV) in $100$ μL of $0.5$ M H<sub>2</sub>SO<sub>4</sub> in water. CV was cycled from $-1.4$ to $+0.4$ V relative to Ag pseudo-reference with $10$ mV step and $300$ mV · sec<sup>-1</sup> scan rate repeatedly until the reduction peak surpassed $-0.5$ mA. Electrodes were then rinsed with DI water and dried with N<sub>2</sub> prior to DNA functionalization.

All DNA sequences used during electrode preparation include a 5' C6-disulfide modification. Before exposure to the electrode, the DNA was first reduced with 1,000-fold molar excess of TCEP for $1$ h to expose reactive thiols.

Electrodes were then incubated for $\geq 5$ hours at room temperature with $500$ nM disulfide-cleaved DNA diluted into immobilization buffer ($1\times$ PBS + $2$ M NaCl) at a volume sufficient to cover the working electrode completely ($\geq 20$ μL). This incubation was conducted in a humidified enclosure, with water added to the base of an empty pipette tip box and electrodes positioned on the elevated tray to maintain a humid environment during incubations. After 5 hour DNA incubation, electrodes were rinsed with 1× PBS and dried gently with N<sub>2</sub>. 100 μL of 5 mM 6-Mercapto-1-Hexanol (MCH) in 1× PBS was subsequently added to each electrode and incubated overnight in the humidified enclosure. Electrodes were again rinsed with 1× PBS, dried gently with N<sub>2</sub>, placed into 100 μL of test buffer and equilibrated for at least 1 h prior to electrochemical testing.

When double-stranded 12-mer duplexes were tested, the electrodes were prepared with the single-stranded disulfide cleaved DNA and a subsequent MCH SAM backfill as described above. The electrode-tethered DNA layers were subsequently hybridized in 100 μL of 1 μM solution of the respective reverse complement in the test buffer for 2 h. The electrode was then washed with 1× PBS and reequilibrated into the test buffer prior to electrochemical testing.

##### *Bio-layer interferometry*

All BLI measurements were conducted on a Gator Plus BLI Instrument using Small Molecule Analysis Probes (SMAP) (Gator Bio, Palo Alto, CA). All BLI experiments utilized the aforementioned Tris-based test buffer used for electrochemical testing. BLI protocols were composed of a $\geq 10$ min buffer equilibration step followed by 120 s of baseline in buffer, 60 s of loading in 500 nM biotinylated probe solution, 240 s of post-loading baseline in buffer, 450 s of association in target solution, and 600 s of dissociation in buffer. All BLI data presented herein are reference-subtracted against a reference probe that was not exposed to DNA during the loading step but was exposed to an identical target solution. The data are further referenced against reference-subtracted response to a zero-target condition. This subtraction ensures the reported curves reflect DNA-aminoglycoside interactions as opposed to probe matrix-aminoglycoside interactions. Presented data are representative.

##### *Circular dichroism*

All circular dichroism measurements were conducted on a Jasco J-815 CD spectropolarimeter (Jasco, Oklahoma City, OK). Prior to measurement, the system was purged with N<sub>2</sub> gas for 10 min. All measurements were conducted with a 150 μL sample volume in a Micro Hellma absorption cuvette with a 1mm path length (Hellma GmbH & Co. KG, Müllheim, Germany). Between measurements, the cuvette was rinsed thoroughly with ethanol and DI water using a side arm flash with a P65 Series Cuvette Washer attachment (Sigma Aldrich C1295) under vacuum. Spectral measurements were taken at room temperature from 320 nm to 220 nm with standard sensitivity and a 1-s digital integration time (DIT) in continuous scanning mode at 50 nm · min<sup>-1</sup>. Three spectra were collected per sample and averaged to produce plots. After averaging, data was smoothed using a third order butterworth filter with a cutoff at 12% of the Nyquist frequency and converted from millidegrees (mdeg) to ellipticity ($\Delta\epsilon$) based on the number of bases in the sequence, the concentration of DNA in the cell, and cuvette path length as per: $CD_{(\Delta\epsilon)} = \frac{CD(mdeg)}{32980 * n_{bases} * [DNA] * path\ length\ (cm)}$. In addition to measurement of CD signal, high tension voltage (HT) was also tracked throughout experiments; these values were below 450V in all reported CD data for aminoglycosides and for the glucose positive control. For the vancomycin positive control, HT signal was higher due to strong absorption by vancomycin near 220nm; the maximum HT measured for this positive control was below 600V. All plotted reference spectra are reference-subtracted averages against buffer only conditions. All plotted spectra with targets are reference-subtracted averages against target in buffer conditions. Semi-transparent patches indicate error-propagated single standard deviations.

For comparing circular dichroism measurements across different DNA sequences, we use a relative structure shift metric (**Table S1**). This metric is calculated by comparing four pre-averaged and pre-smoothed measurements: DNA without target, DNA with target, buffer only, and buffer with target spectra. We correct the DNA without target spectra by subtracting the buffer only spectra from it. We then correct the DNA with target spectra by subtracting the buffer with target spectra from it. We then calculate the integral of the absolute value of the difference between corrected DNA with target and corrected DNA without target spectra. Finally, we normalize this integrated difference by the integral of the absolute value of the corrected DNA without target spectra. The resulting metric therefore compares differences in ellipticity normalized against a measure of total baseline ellipticity.

##### *In-situ electrode imaging under voltage*

Jumper wires were soldered to Metrohm 220BT electrodes and then cleaned and functionalized as described above with Cy3- or Cy3B-labeled disulfide-cleaved DNA. Electrodes were imaged facing down toward the objective under a custom, transparent Grace Bio-Labs (Bend, Oregon) plastic chamber filled with the testing solution positioned on a custom mounting slide. Electrodes were interfaced to a emStat4s LR potentiostat via the soldered jumper wires. Imaging was performed on an inverted Nikon Eclipse Ti2 microscope using a 4x Plan Apo objective (Nikon MRD70040). The effective pixel size was 5.5 $\mu\text{m}$. A 532nm laser and 561nm laser (LUN-F XL 532/561/640 Laser Combiner) were used for excitation of Cy3 and Cy3B constructs, respectively. Cy3 and Cy3B emission fluorescence was filtered with ET585/65m and ET600/50 (Chroma Technology) filters, respectively. All movies were recorded onto a 600 × 600 pixel region of a back-illuminated Scientific CMOS camera (Prime 95B, 1.44 MP, Teledyne Photometrics). The camera and microscope were controlled using the NIS-Elements Advanced Research Software Package. Excitation power of 14 mW 532nm and 47.4mW, 561nm was used throughout imaging. Videos were recorded at an effective 80 frames per second. Nominal exposure setting was 10 ms. The potentiostat was controlled in PStace 5.13 software and used to apply an SWV waveform during imaging. SWV ramped from $-0.45$ to $-0.1$ V with a 1 mV step, 25 mV amplitude after a 1 s pre-conditioning time (voltage held at

0 V) and a 1 s equilibration time (voltage held at $-0.45$ V). To exchange solutions, the chamber was gently removed, excess solution was wicked from the surface with a Kimwipe, and a new solution was allowed to incubate on the electrode as a sessile 100-μL droplet for at least 5 min. The droplet was then wicked from the surface, and a second droplet with 20 μL volume was applied to the electrode, and the chamber was replaced over the electrode. The electrode was then positioned for the next round of imaging.

Images were analyzed using a circular region-of-interest (ROI) in the center of the image (radius 200 pixels). Average fluorescence intensity-time trajectories within the ROI were computed in Python. Reported values include subtraction of a background level computed from imaging an electrode not functionalized with DNA. Baseline fluorescence is determined from the average of background subtracted values during the 0 V pre-conditioning period prior to the application of $-0.45$ V, as determined by the step in fluorescence observed in all computed intensity traces. Presented data are representative examples.

#### *Fluorescence assays*

All in-solution fluorescence measurements were performed on a BioTek Synergy H1 microplate reader using CoStar 96-well, half-area opaque plates (Corning 3993). For positive control conditions using the sequence's unlabeled reverse complement (and reference conditions), an annealing step in our test buffer (3 minutes at $95^{\circ}\text{C}$, stepping down $4^{\circ}\text{C}$ per minute until the sample reached room temperature) was performed prior to plating. When used, the reverse complement was in 1.5-fold molar excess of the labeled test sequence. Estimated concentrations of labeled components during annealing were 1 $\mu\text{M}$ and 3 $\mu\text{M}$ for the labeled aminoglycoside aptamer and hairpin control, respectively. Final concentrations were 50 nM and 150 nM for the labeled aminoglycoside aptamer and hairpin controls, respectively. Final volumes were 80 $\mu\text{L}$ per well. Fluorescence measurements were collected with an excitation wavelength of 590 nm and an emission wavelength of 620 nm. Presented results represent the average and standard deviation of three replicates.

MB fluorescence measurements were performed on the same instrument and plates described for the Alexa Fluor 594 assays with an excitation wavelength of 660 nm and emission wavelength of 695 nm. DNA-MB concentration was fixed at 400 nM final concentration at 80 $\mu\text{L}$ per well. Presented results represent the average and standard deviation of three replicates.

In-solution fluorescence of Cy3-labeled DNA was measured using a filter cube with a 540/25 nm excitation filter, a 590/35 nm emission filter, and 570 nm top mirror. In-solution fluorescence of Cy3B-labeled DNA was measured using an excitation wavelength of 550 nm and an emission wavelength of 580 nm. Final DNA concentrations were 50 nM at 80 $\mu\text{L}$ per well. Presented results represent the average and standard deviation of three replicates.

*pH assay*

pH of buffers were verified using an AE150 Accumet (Fisher Scientific) instrument and probe (13-620-299B). pH-adjusted buffer for **Fig. S3** was prepared by dropwise addition of 1 N NaOH or 1 N HCl solutions.

### SI Figures

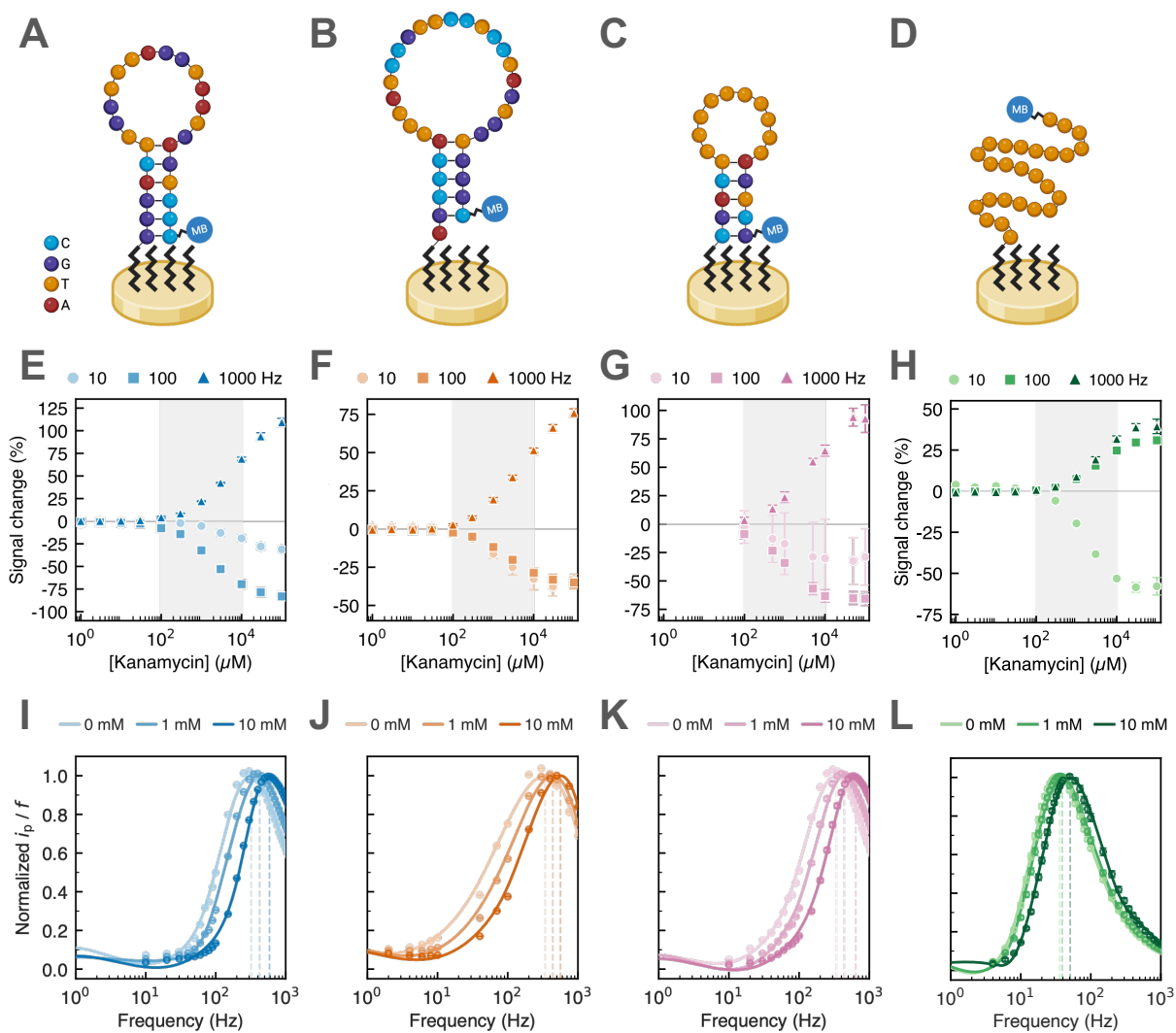

**Figure S1. Kanamycin elicits a large electrochemical signal by increasing electron transfer from multiple different DNA-MB constructs.** (A–D) Illustrative representations of predicted secondary structures for (A) the canonical aminoglycoside aptamer, (B) the IgGE DNA aptamer from [10], (C) a random hairpin sequence, and (D) a poly-T<sub>26</sub> control sequence. (E–H) Square-wave voltammetry (SWV) signal response curves and (I–L) plots of frequency-normalized peak current across frequency for (E, I) the canonical aminoglycoside aptamer, (F, J) the IgGE aptamer, (G, K) the random hairpin control, and (H, L) the poly-T<sub>26</sub> control. All sequences were modified distally with methylene blue (MB). Solid lines in I–L indicate spline fits to frequency-normalized current peaks. Dashed lines show the maxima of the frequency-normalized current peaks, which are used as proxies for ensemble electron transfer rates [34–36].

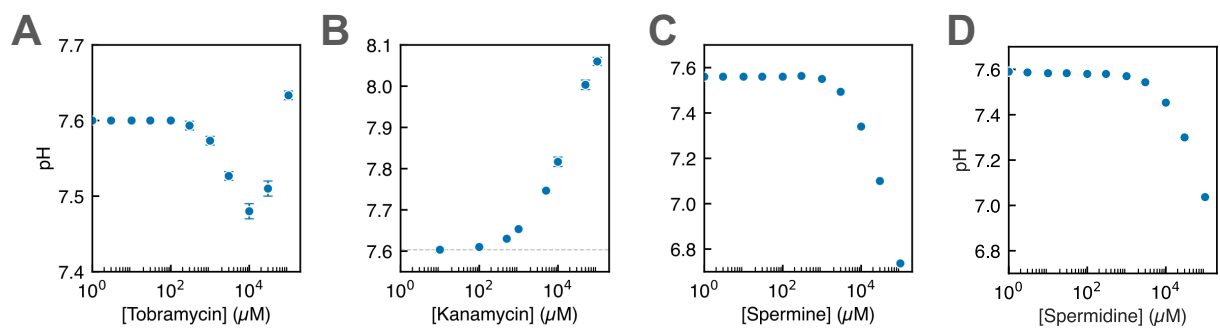

**Figure S2. Buffer pH changes differently in response to aminoglycosides and polyamines.** pH response of our standard testing buffer to increasing concentrations of (A) tobramycin, (B) kanamycin, (C) spermine, and (D) spermidine. Error bars represent one standard deviation of three replicates. Refer to **Figure S3C** for impact of kanamycin on pH-adjusted buffer.

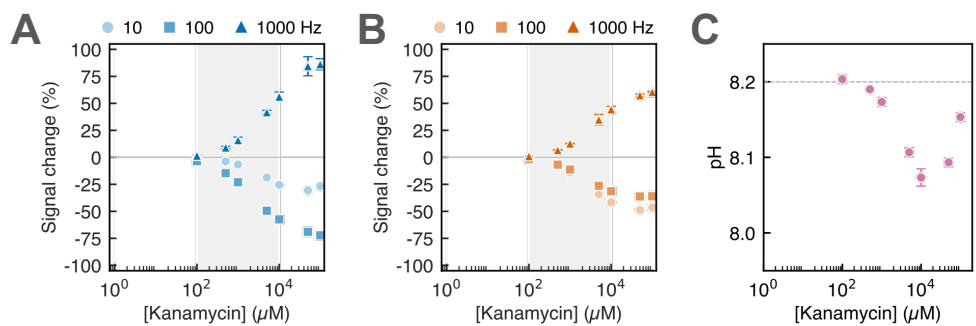

**Figure S3. pH effects do not explain the sequence-agnostic electrochemical response of MB-DNA to aminoglycosides.** SWV response of (A) the canonical aminoglycoside aptamer and (B) the IgGE aptamer control to kanamycin in a 20 mM Tris, 100 mM NaCl, 5 mM MgCl<sub>2</sub> buffer pre-adjusted to approximately pH 8.2. (C) The pH of the pre-adjusted buffer shows minimal pH variation with kanamycin titration. Error bars represent one standard deviation of three replicates. For comparison to standard buffer response to kanamycin titration, see **Figure S2B**.

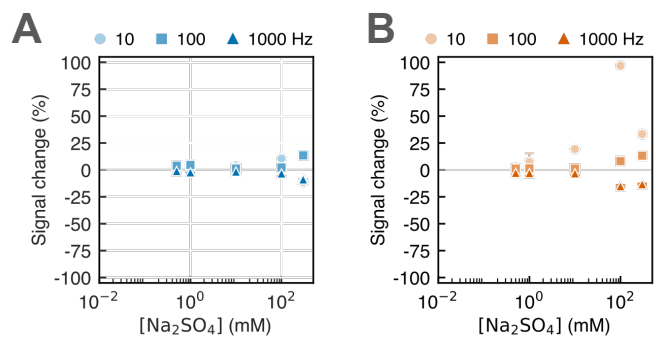

**Figure S4. Ionic strength does not explain the sequence-agnostic electrochemical response of MB-DNA to aminoglycosides.**

SWV responses of (A) the canonical aminoglycoside aptamer and (B) the IgGE aptamer control to sodium sulfate titrations up to concentrations that are in considerable excess of those encountered in aminoglycoside experiments. The 10 Hz IgGE aptamer response to 100 mM sodium sulfate is notable but opposite to the responses to tobramycin and kanamycin. Error bars represent one standard deviation of three replicates.

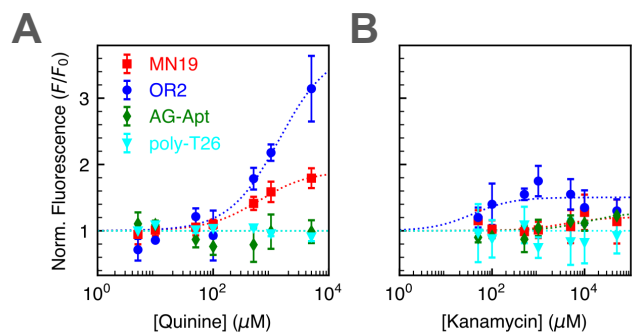

**Figure S5. Fluorescence recovery of MB-labeled DNA rules out redox reporter-ligand competition for aminoglycoside EAB signaling.** (A) Redox reporter-ligand competition is known to occur with cocaine/quinine-binding sequences MN19 and OR2 based on recovery of MB fluorescence upon titration of target, similar to the report in [23]. In contrast, the canonical aminoglycoside aptamer and control poly-T<sub>26</sub> do not indicate MB fluorescence recovery in response to quinine. (B) MB fluorescence recovery for the same sequences in response to kanamycin titration shows no evidence of significant fluorescence recovery. Error bars represent one standard deviation of three replicates.

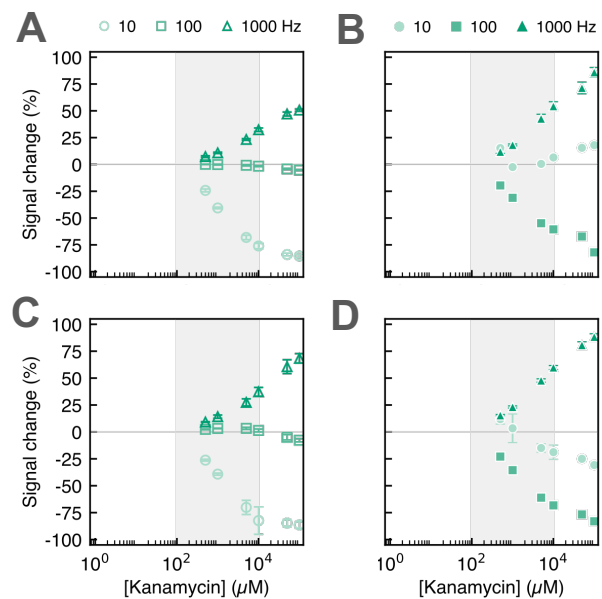

**Figure S6. Two variants of proximally MB-labeled 12-mer ssDNA and dsDNA with random sequence respond to kanamycin titrations.** Response to kanamycin titration of (A) a proximally MB-labeled 12-mer ssDNA (Variant 1) alone or (B) hybridized to its reverse complement, as well as (C) an alternative proximally MB-labeled 12-mer ssDNA (Variant 2) alone or (D) hybridized to its reverse complement. Error bars represent one standard deviation of three replicates.

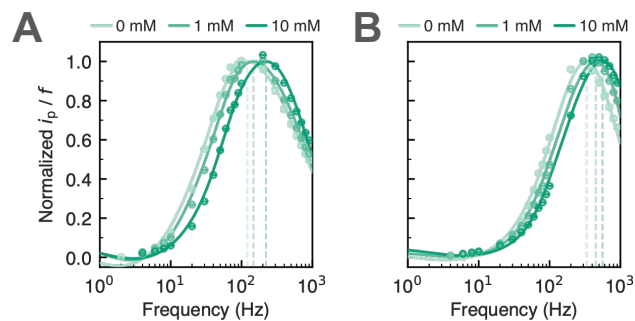

**Figure S7. Mirčeski analysis demonstrates differences in electron transfer rate for 12-mer ssDNA and dsDNA.** Mirčeski analysis of (A) unhybridized Variang 1 ssDNA and (B) the hybridized Variant 1 dsDNA response to a titration of tobramycin. Hybridization increases baseline apparent transfer rate nearly three-fold. Error bars represent one standard error on the mean of three replicates.

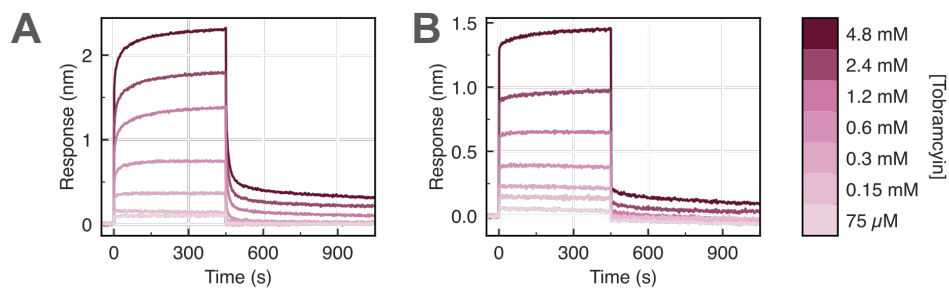

**Figure S8. BLI shows promiscuous association of tobramycin to additional control DNA strands.** BLI demonstrates association and dissociation of tobramycin for (A) a scrambled version of the legacy aminoglycoside sequence and (B) a hairpin control (alt-SLP-T10). Association occurs from $t = 0$ to $t = 450$ s. Dissociation occurs for $t > 450$ s.

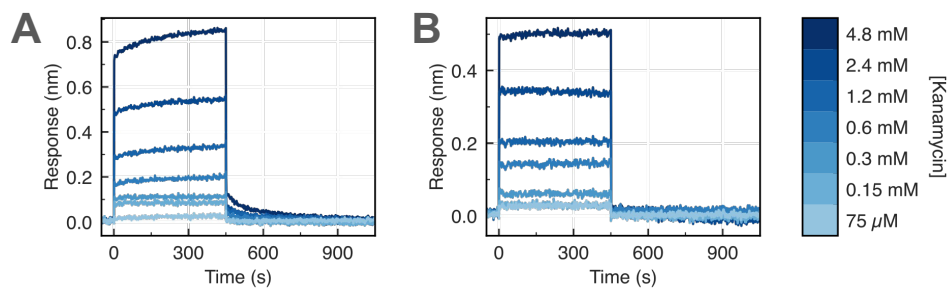

**Figure S9. BLI shows promiscuous association of kanamycin to DNA.** BLI demonstrates association and dissociation of kanamycin for (A) the legacy aminoglycoside sequence and (B) a hairpin control (alt-SLP-T10). Association occurs from $t = 0$ to $t = 450$ s. Dissociation occurs for $t > 450$ s.

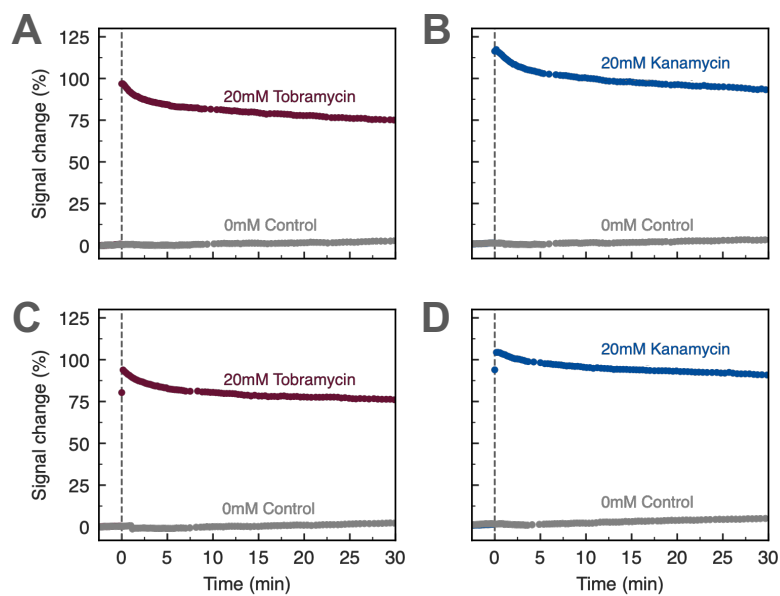

**Figure S10. Time-resolved SWV responses corroborate rapid non-specific aminoglycoside association with DNA.** Representative SWV responses of (A, B) the canonical aminoglycoside aptamer or (C, D) a hairpin control to a 20 μL spike-in of 100 mM (A, C) tobramycin or (B, D) kanamycin (versus buffer spike-in) into 80 μL of buffer (final 20 mM aminoglycoside concentration) at $t = 0$. SWV frequency = 1,200 Hz. A 3-s wait step was deployed between each SWV interrogation. Signal decrease after initial response suggests diffusion of the spike bolus into the larger droplet volume.

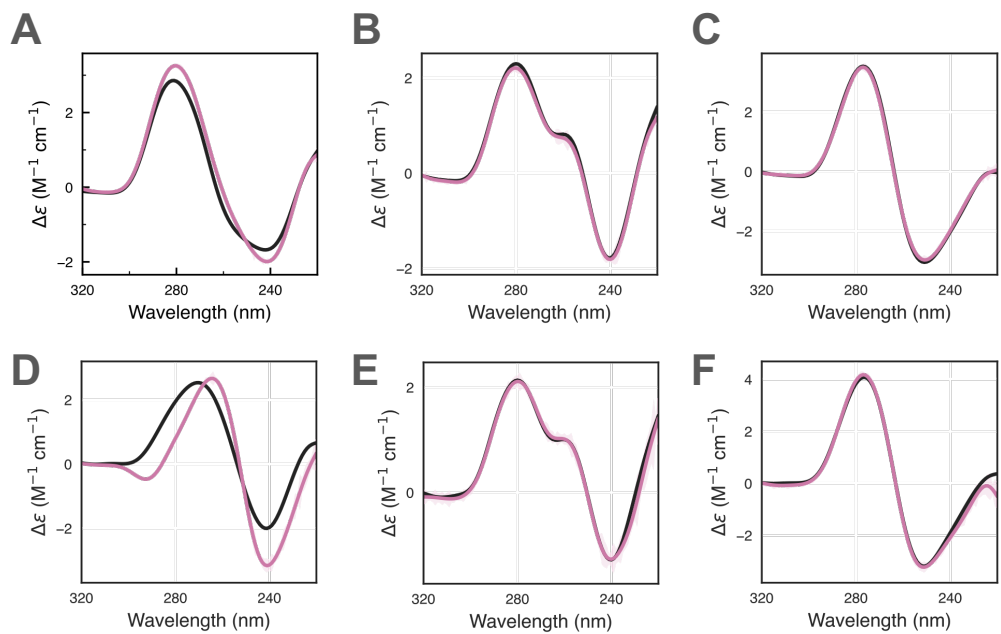

**Figure S11. Structure-switching aptamers exhibit CD spectral shifts.** (A) CD data for a glucose-sensing beacon with known structure-switching behavior. Spectra are shown in the absence (black) and presence (colored) of 100 mM glucose. This sequence was mutated to destabilize its stem region in the target free state. Upon introduction of glucose, the stem closes, increasing the dsDNA signature detectable via CD. (B, C) Negative control responses to 100 mM glucose for (B) the canonical aminoglycoside aptamer and (C) poly-T<sub>26</sub>. (D) CD data for a vancomycin-sensing aptamer sequence with known structure-switching properties in the absence (black) and presence (colored) of 100 μM vancomycin. (E, F) Negative control responses to 100 μM vancomycin for (E) the canonical aminoglycoside aptamer and (F) poly-T<sub>26</sub>. Semi-transparent patches represent error-propagated single standard deviations. Quantitative relative shifts are available in **Table S1**.

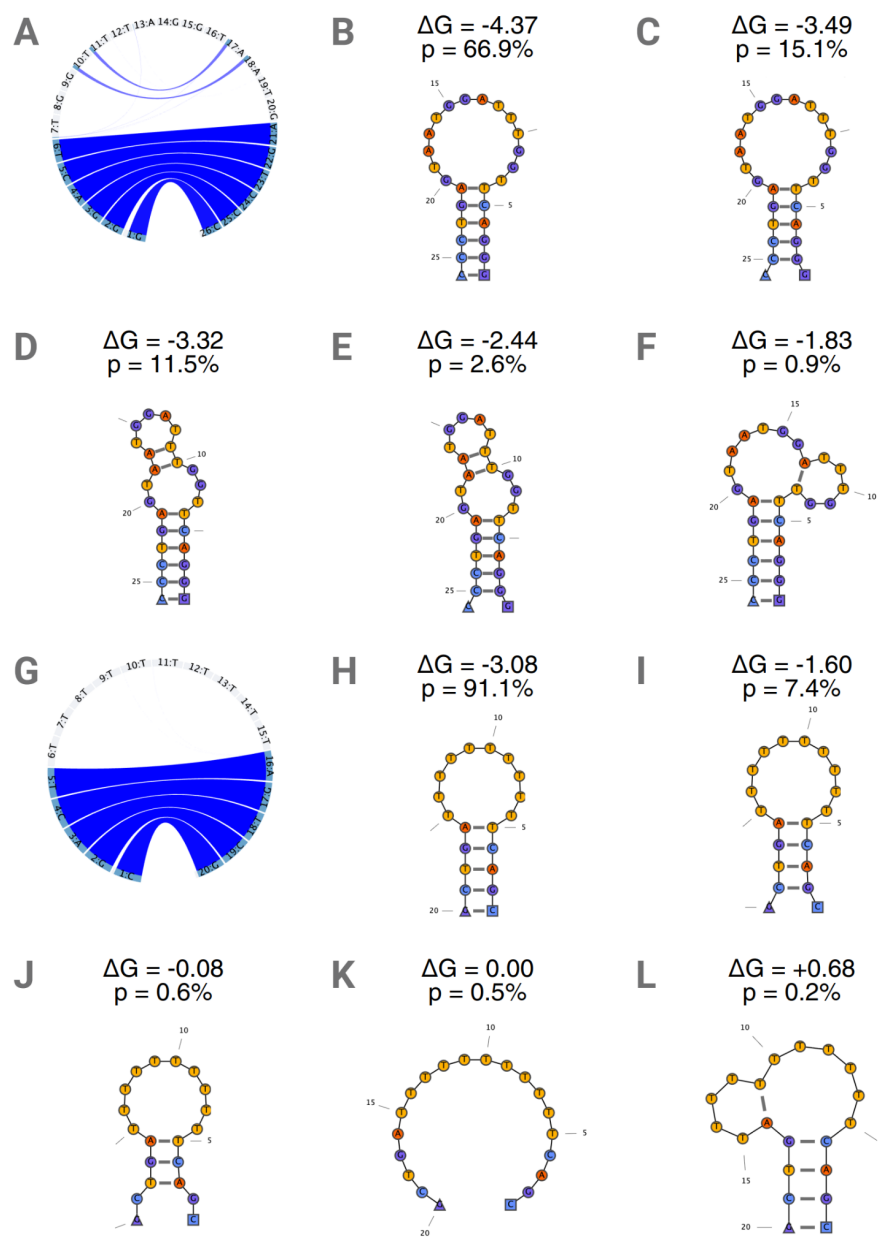

**Figure S12. Predicted unbound secondary structures of the legacy aminoglycoside and control hairpin.** NUPACK predicts a predominance of hairpin secondary structures for (A–F) the aminoglycoside aptamer sequence and (G–L) a control stem-loop probe (alt-SLP-T10) in the absence of target under conditions matching our test buffer. (A, G) are chord diagrams representing probability of base pairings across the 500 most probable secondary structures as widths of connecting chords. The top five predicted secondary structures in descending order are shown for the canonical aminoglycoside aptamer and hairpin control in (B–F) and (H–L), respectively, with annotated $\Delta G$ and probability values. All $\Delta G$ values are reported in units of $\text{kcal} \cdot \text{mol}^{-1}$. The top five structures represent 97% and 99.8% of the predicted populations of the aminoglycoside aptamer and stem-loop probe, respectively [52].

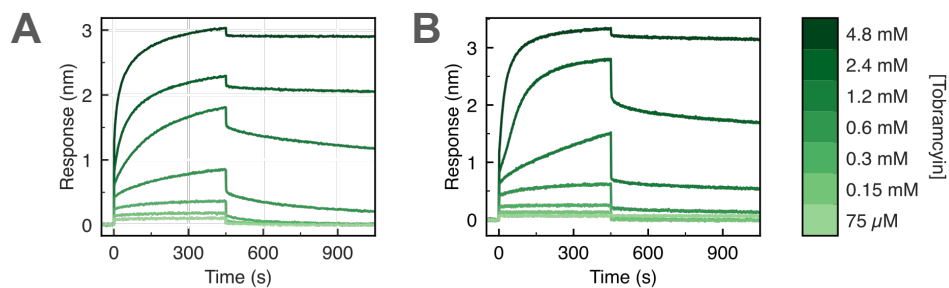

**Figure S13. BLI demonstrates promiscuous spermine association to DNA and limited dissociation.** BLI demonstrates association and limited dissociation of spermine to (A) the canonical aminoglycoside aptamer and (B) a hairpin control. Association occurs from $t = 0$ to $t = 450$ s. Dissociation occurs for $t > 450$ s.

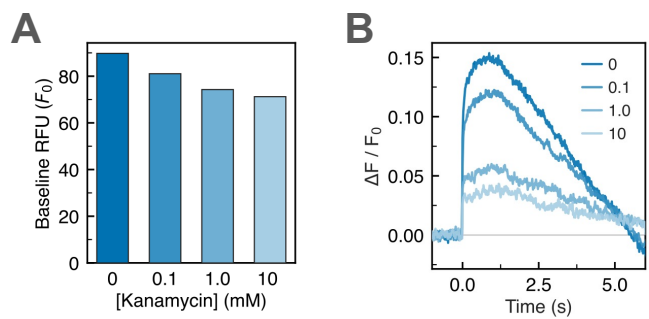

**Figure S14. Fluorescence analysis of electrodes modified with Cy3-labeled aminoglycoside aptamer exposed to kanamycin.** (A) The baseline fluorescence of the Cy3-labeled aptamer decreases with increasing kanamycin concentration in a 20 mM Tris, 120 mM NaCl, 5 mM KCl, 1 mM $\text{MgCl}_2$, 1 mM $\text{CaCl}_2$ buffer. (B) Representative mean fluorescence intensity of 3'-Cy3-labeled aminoglycoside aptamer immobilized onto an electrode and subjected to 100 Hz SWV interrogation with increasing kanamycin concentration in the same buffer. All concentrations are millimolar.

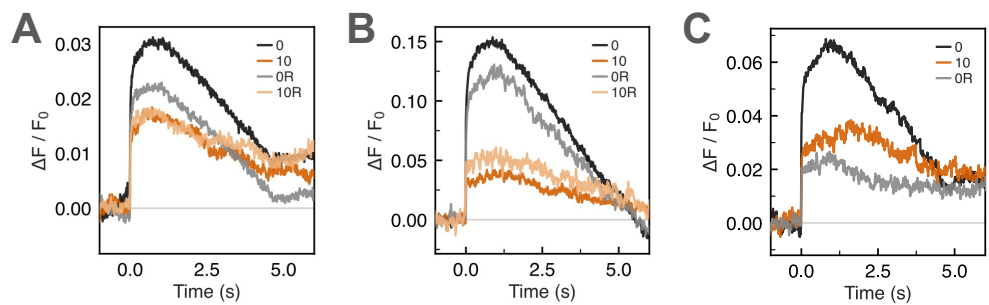

**Figure S15. Reversibility of fluorescence modulation changes.** Representative mean fluorescence intensity of 3'-Cy3-labeled aminoglycoside aptamer immobilized onto electrodes and subjected to 100 Hz SWV interrogation while exposed to 10 mM (A) tobramycin, (B) kanamycin, or (C) spermine, alternated with buffer washes. For tobramycin and kanamycin, some hysteresis is observed likely due to incomplete washing and/or photobleaching. For spermine, poor reversibility is observed. Kanamycin condition taken in 20 mM Tris, 120 mM NaCl, 5 mM KCl, 1 mM MgCl<sub>2</sub>, 1 mM CaCl<sub>2</sub> buffer. All concentrations are indicated in units of millimolar, with 'R' indicating a repeated condition.

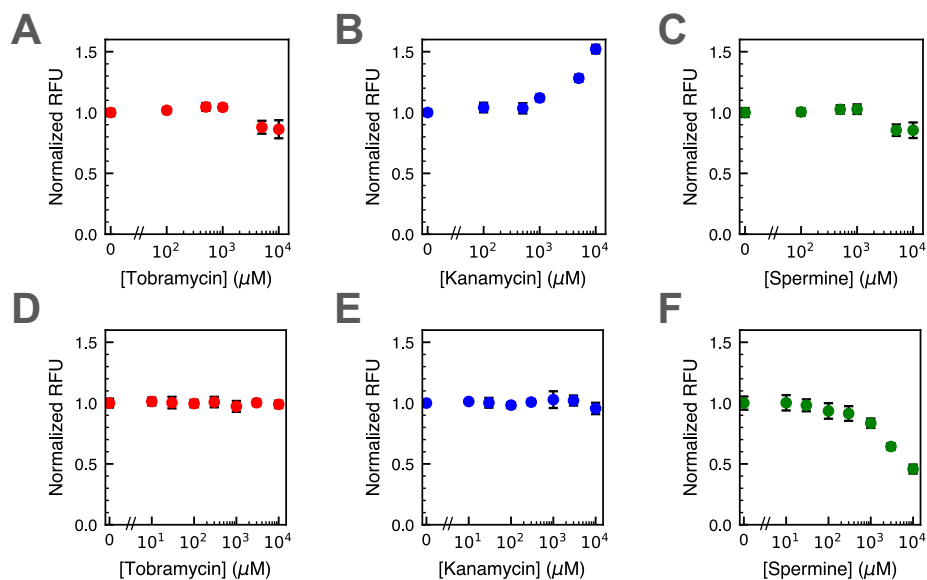

**Figure S16. In-solution fluorescence controls for Cy3- and Cy3B-labeled DNA.** (A–C) Normalized fluorescence intensities of Cy3-labeled aminoglycoside aptamer in testing buffer solution against increasing concentrations of (A) tobramycin, (B) kanamycin, and (C) spermine. (D–F) Normalized fluorescence intensities of Cy3B-labeled poly-T<sub>26</sub> in testing buffer solution against increasing concentrations of (D) tobramycin, (E) kanamycin, and (F) spermine. Error bars represent the standard deviation of three replicates. These controls indicate that fluorescence quenching on electrode cannot be explained by direct effects of the titrated targets on either fluorophore except for spermine against the poly-T<sub>26</sub>.

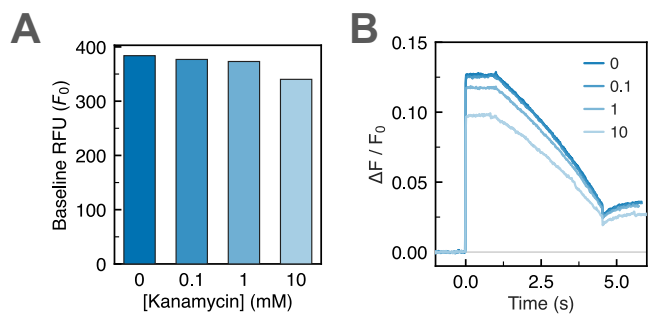

**Figure S17. Muted fluorescence quenching and modulation reduction for the poly-T<sub>26</sub>-Cy3B in response to kanamycin.** (A) Baseline fluorescence of 3'-Cy3-labeled poly-T<sub>26</sub> against increasing concentrations of kanamycin. (B) Representative mean fluorescence intensity of 3'-Cy3-labeled poly-T<sub>26</sub> under 100 Hz SWV interrogation in standard testing buffer in response to a titration of kanamycin. The muted result mirrors the reduced sensitivity of this sequence to kanamycin in electrochemical assays. All concentrations are millimolar.

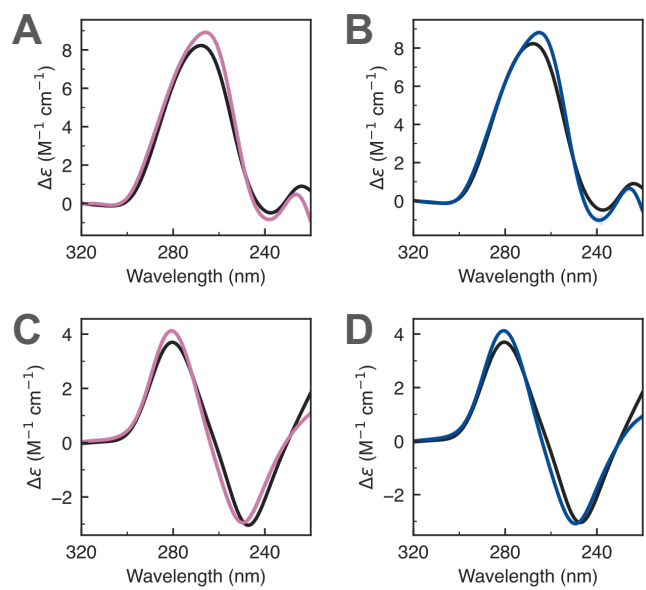

**Figure S18. CD spectra for the 2'-OMe version of the legacy aminoglycoside aptamer and a newly-identified kanamycin aptamer.** (A, B) CD spectra for the distally MB-labeled full 2'-OMe homolog of canonical aminoglycoside aptamer in the absence (black) or presence (color) of 100 mM (A) tobramycin and (B) kanamycin. (C, D) CD spectra of distally MB-labeled kanamycin aptamer KAN 8-1 in the absence (black) or presence (colored) of 100 mM (C) tobramycin and (D) kanamycin. Semi-transparent patches represent error-propagated single standard deviations. Quantitative comparisons available in **Table S1**.

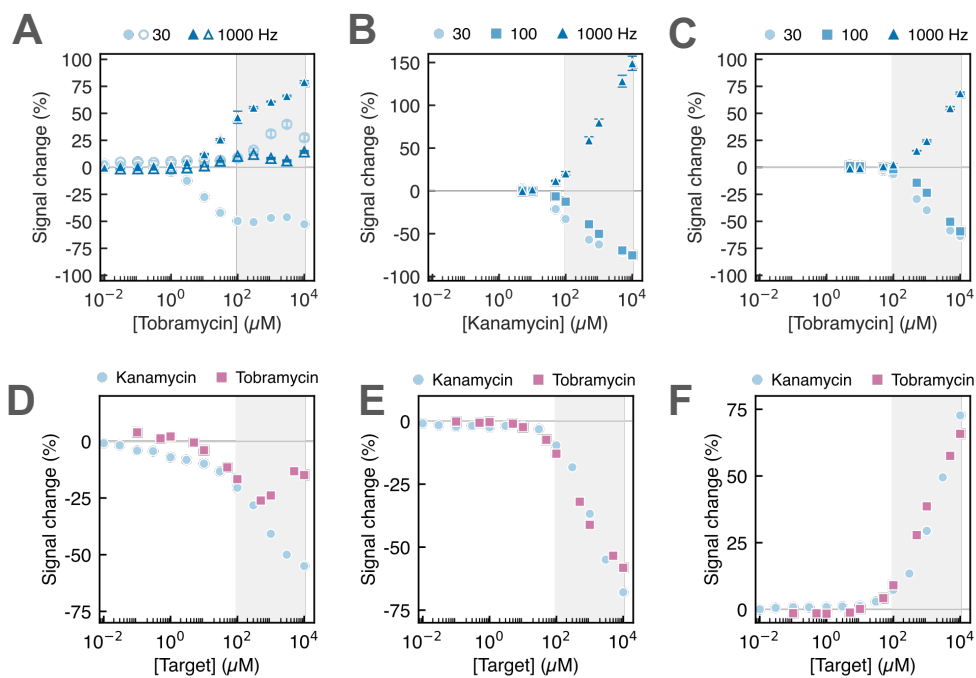

**Figure S19. Electrochemical evaluation of EABs based on recently published aminoglycoside aptamers.** (A) SWV peak current responses of a distally MB-labeled 2'-OMe version of the legacy aminoglycoside aptamer (solid markers) demonstrate enhanced affinity of this construct toward tobramycin as compared to its proximally MB-labeled counterpart (open markers). (B) In contrast, the SWV response to kanamycin of a distally MB-labeled 2'-OMe version of the legacy aminoglycoside aptamer does not show sensitivity beyond charge-screening effects observed for its DNA counterpart. (C) A control 2'-OMe stem-loop probe does not show sensitivity beyond charge-screening effects in response to tobramycin. These results suggest that the 2'-OMe aminoglycoside EAB improves affinity and specificity of the sensor to tobramycin via a mechanism separate from charge-screening. (D) KAN 8-1 SWV peak current response at 30 Hz shows partially-enhanced sensitivity to kanamycin relative to tobramycin at concentrations < 100 $\mu\text{M}$. KAN 8-1 SWV responses at (E) 100 Hz and (F) 1,000 Hz do not distinguish between kanamycin and tobramycin, and respond only in the regime where charge-screening effects are also observed for control sequences. Error bars represent standard deviations.

**Table S1. CD relative structure-shift metrics (%)**

| **Sequence** | **vs. 100 mM tobramycin** | **vs. 100 mM kanamycin** | **vs. 100 mM glucose\*** | **vs. 100 μM vancomycin\*\*** |
| --- | --- | --- | --- | --- |
| **Unmod-AG-Apt** | 8.193 | 7.382 | 4.446 | 5.876 |
| **SLP-T10-Prox (unlabeled)** | 5.959 | 4.982 | - | - |
| **Unmod-Poly-T<sub>26</sub>** | 7.038 | 3.401 | 3.757 | 7.125 |
| **GLU-01** | - | - | 18.152 | - |
| **Unmod KAN 8-1** | 15.284 | 17.738 | - | - |
| **AG-Apt-XNA (unlabeled)** | 14.574 | 11.509 | - | - |
| **Unmod-Vanco-Apt** | - | - | - | 64.722 |

All relevant aminoglycoside spectra conducted in 20 mM Tris, 100 mM NaCl, 5 mM MgCl<sub>2</sub> testing buffer.

\*All relevant glucose spectra conducted in 1× HEPES + 1 M NaCl + 10 mM MgCl<sub>2</sub> + 5 mM KCl buffer.

\*\* All relevant vancomycin spectra conducted in 1× PBS + 2 mM MgCl<sub>2</sub> buffer.

- Data not collected.

**Table S2. DNA sequences**

| ID | Description | Sequence | Source | Used In |
| --- | --- | --- | --- | --- |
| AG-Apt | Thiolated canonical aminoglycoside aptamer with 3' amino C6 group for MB or Cy3 labeling | /5ThioMC6-D/GGGACTTGGTTTAGGTAATGAGTCCC/3AmMO/ | IDT | 1E, 1I, 2B, 6A, 6B, 6C, 6D, 6E, 6F, 7B, 7C, 7D, 7E, S1E, S1I, S3A, S4A, S10A, S10B, S14, S15, S16, S19 |
| IgGE-Apt | Thiolated IgGE-binding aptamer with 3' amino C6 group for MB labeling | /5ThioMC6-D/AGCCCATTTATCCGTTCCTCCTAGTGGTGGGC/3AmMO/ | In-house | 1F, 1J, S1F, S1J, S3B, S4B |
| SLP-T10 | Thiolated stem loop probe variant 1 with 3' amino C6 group for MB labeling | /5ThioMC6-D/GCGAATTTTTTTTTTTTCGC/3AmMO/ | In-house | 1G, 1K, 2C, 6C, 6D, 6E, 6F, S1G, S1K, S10C, S10D |
| Poly-T<sub>26</sub> | Thiolated poly-T<sub>26</sub> with 3' amino C6 group for MB or Cy3B labeling | /5ThioMC6-D/TTTTTTTTTTTTTTTTTTTTTTT/3AmMO/ | IDT | 1H, 1L, 6C, 6D, 6E, 6F, 7F, 7G S1H, S1L, S16, S17, S19 |
| AG-Apt-Prox | Thiolated canonical aminoglycoside aptamer with proximal Unilink amino group for MB labeling | /5ThioMC6-D//iUniAmM/GGGACTTGGTTTAGGTAATGAGTCCC | IDT | 2B |
| SLP-T10-Prox | Thiolated stem loop probe variant 1 with proximal Unilink amino group for MB labeling or unlabeled for CD | /5ThioMC6-D//iUniAmM/GCGAATTTTTTTTTTTTCGC | IDT | 2C, 5B, 5F, S11C, S11F |
| Alt-SLP-T10 | Thiolated stem loop probe variant 2 with 3' amino C6 group for MB labeling | /5ThioMC6-D/CGACTTTTTTTTTTTAGTCG/3AmMO/ | IDT | 2D |
| Alt-SLP-T10-Prox | Thiolated stem loop probe variant 2 with proximal Unilink amino group for MB labeling | /5ThioMC6-D/iUniAmM/CGACTTTTTTTTTTTAGTCG | IDT | 2D |
| R-12m | Thiolated random 12-mer variant 1 with proximal Unilink amino group for MB labeling | /5ThioMC6-D//iUniAmM/GGCATTGCTACA | IDT | 2E, 2F, S6A, S6B, S7A, S7B |
| R-12m-Comp | Reverse complement to R-12m | TGTAGCAATGCC | IDT | 2F, S6B, S7B |
| Bio-AG-Apt | 5' biotinylated canonical aminoglycoside aptamer | /5BiotinTEG/GGGACTTGGTTTAGGTAATGAGTCCC | IDT | 4A, S9A, S13A |
| Bio-IgGE-Apt | 5' biotinylated IgGE aptamer | /Biotin/AGCCCATTTATCCGTTCCTCCTAGTGGTGGGC | In-house | 4B |
| Bio-SLP-T10 | 5' biotinylated stem loop probe variant 1 | /Biotin/GCGAATTTTTTTTTTTTCGC | In-house | 4C |
| Bio-Poly-T<sub>26</sub> | 5' biotinylated poly T<sub>26</sub> | /Biotin/TTTTTTTTTTTTTTTTTTTTTTTTTT | In-house | 4D |
| Unmod-AG-Apt | Unmodified canonical aminoglycoside aptamer | GGGACTTGGTTTAGGTAATGAGTCCC | IDT | 5A, 5E, S11B, S11E |
| Unmod-Poly-T<sub>26</sub> | Unmodified poly T<sub>26</sub> | TTTTTTTTTTTTTTTTTTTTTTTTTT | IDT | 5C, 5G, S11C, S11F |
| FRET-AG-Apt | 5' BHQ-2 canonical aminoglycoside aptamer with 3' amino C6 for AF594 labeling | /5' BHQ2/GGGACTTGGTTTAGGTAATGAGTCCC/3' AminoC6/ | In-house | 5D, 5H |
| AG-Apt-Comp | Reverse complement to canonical aminoglycoside aptamer | GGGACTCATTACCTAAACCAAGTCCC | In-house | 5D, 5H |
| FRET-Alt-SLP-T10 | 5' amino C6 stem loop probe variant 2 with 3' BHQ-2 modification. Subsequent AF594 labeling. | /5'DMS(o)MT-Amino/CGACTTTTTTTTTTAGTCG/3'BHQ2/ | In-house | 5D, 5H |
| Alt-SLP-T10-Comp | Reverse complement to stem loop probe variant 2 | CGACTAAAAAAAAAAGTCG | In-house | 5D, 5H |
| R-12m-2 | Thiolated random 12-mer variant 2 with proximal Unilink amino group for MB labeling | /5ThioMC6-D/iUniAmM/ACTGATCGTACA | IDT | S6C, S6D |
| R-12m-2-Comp | Reverse complement to R-12m-2 | TGTACGATCAGT | IDT | S6D |
| Bio-AG-Scram | 5' biotinylated scrambled aminoglycoside aptamer sequence | /Biotin/ATTCTGCTGTGATAGCTGGATGCGGA | In-house | S8A |
| Bio-Alt-SLP-T10 | 5' biotinylated stem loop probe variant 2 | /Biotin/CGACTTTTTTTTTTAGTCG | In-house | S8B, S9B, S13B |
| GLU-01 | Glucose binding aptamer beacon for positive control CD | /Biotin//Sp18//dabcyldT/TGGACCACCGTGTGTGTTGCTCTGT AACAGTGTCCATTGTCGTCCCT/3'AminoC3/ | In-house | S11A |
| Unmo-Vanco-Apt | Unmodified Vanco EAB sequence for positive control CD | CGAGGGTACCGCAATAGTACTTATTGTTCGCCTATTGTGGGTCGG | IDT | S11D |
| Unmod KAN 8-1 | Unmodified KAN 8-1 for CD | GACGACGCAGTCGGCTTTTGC GGGA AAAAGGTCGGAGTCGTC | IDT | S18C, S18D |
| AG-Apt-XNA | 5' thiolated, 2'-OMe homolog of canonical aminoglycoside aptamer with 3' amino C3 group for MB labeling. Unlabeled for CD. | /5ThioMC6-D/mGmGmGmAmCmUmUmGmGmUmUmAmGmGmUmAmAmUmGmAmGmUmCmCmC/3' AminoC6/ | In-house | S18A, S18B, S19A, S19B |
| SLP-T10-XNA | 5' thiolated 2'-OMe homolog of stem loop probe variant 1 | /5ThioMC6-D/mGmCmGmAmAmUmUmUmUmUmUmUmUmUmUmUmCmGmC/3AmMO/ | In-house | S19C |
| KAN 8-1 EAB | 5' thiolated KAN 8-1 with 3' amino C6 group for MB labeling | /5ThioMC6-D/GACGACGCAGTCGGCTTTTGC GGGAAAAAGGTCGGAGTCGTC/3' AminoC6/ | In-house | S19D, S19E, S19F |
| MN19 | 5' thiolated MN19 with 3' amino C6 for MB labeling | /5ThioMC6-D/GACAAGGAAAATCCTTCAACGAAGTGGGTC/3' AminoC6/ | In-house | S5 |
| OR2 | 5' thiolated OR2 with 3' amino C6 for MB labeling | /5ThioMC6-D/GACAGGGGGAACCCCTCAACGAAGTGGGTC/3' AminoC6/ | In-house | S5 |

Modified IDT-synthesized sequences are as per IDT modification codes.

In-house synthesized sequences utilize those modifications listed in the materials information.
